# Physiological role and mechanisms of action for a long noncoding haplotype region

**DOI:** 10.1101/2024.05.15.594413

**Authors:** Hong Xue, Manoj K. Mishra, Yong Liu, Pengyuan Liu, Michael Grzybowski, Rajan Pandey, Kristie Usa, Mark A. Vanden Avond, Niharika Bala, Abdel A. Alli, Allen W. Cowley, Qiongzi Qiu, Andrew S. Greene, Sridhar Rao, Caitlin C. O’Meara, Aron M. Geurts, Mingyu Liang

## Abstract

Most common sequence variants associated with human traits are in noncoding regions of the genome, form haplotypes with other noncoding variants, and exhibit small effect sizes in the general population. Determining the physiological roles and mechanisms of action for these noncoding variants, particularly large haplotypes containing multiple variants, is both critical and challenging. To address this challenge, we developed an approach that integrates physiological studies in genetically engineered and phenotypically permissive animal models, precise editing of large haplotypes in human induced pluripotent stem cells (hiPSCs), and targeted chromatin conformation analysis. We applied this approach to examine the blood pressure associated rs1173771 locus, which includes a haplotype containing 11 single nucleotide polymorphisms (SNPs) spanning 17.4 kbp. Deleting the orthologous noncoding region in the genome of the Dahl salt-sensitive rat attenuated the salt-induced increase in systolic blood pressure by nearly 10 mmHg. This attenuation of hypertension appeared to be mediated by upregulation of the adjacent gene *Npr3* (natriuretic peptide receptor 3) in arteries, enhancing vasodilation. The blood pressure-elevating and -lowering haplotypes were precisely reconstituted in hiPSCs using an efficient, two-step genome editing technique. The blood pressure-elevating haplotype decreased NPR3 expression in endothelial cells and vascular smooth muscle cells derived from the edited, isogenic hiPSCs. The influence of the haplotype was partially recapitulated by the sentinel SNP rs1173771. Additionally, the blood pressure-elevating haplotype showed significantly greater chromatin interactions with the *NPR3* promoter region. This study illustrates the feasibility of ascertaining the physiological roles and mechanisms of action for large noncoding haplotypes. Our efficient, integrated, and targeted approach can be applied to investigate other noncoding variants.

## INTRODUCTION

Most single nucleotide polymorphisms (SNPs) associated with human traits are in noncoding regions of the genome.^1^ Understanding how these noncoding sequence variants influence the traits is one of the greatest and most pressing challenges in human genetics.^2–4^ High-throughput approaches such as mapping expression quantitative trait loci (eQTL), massively parallel reporter assays, CRISPR screens, and epigenomic profiling offer valuable insights into the potential effector genes of noncoding variants and possible mechanisms involved.^5–8^ However, direct and targeted testing is essential to ascertain the effect of noncoding loci on in vivo phenotypes and to elucidate the mechanisms in a cellular context relevant to a trait.

Such direct testing is largely absent, contributing to a critical gap between genetic discoveries and physiological and pathophysiological understanding. The reported effect sizes of these loci are often so small that they cannot be detected in animal model studies. Moreover, the limited sequence conservation of noncoding genomic regions across species complicates the identification of specific nucleotides corresponding to human SNPs in model organisms. Additionally, many noncoding variants are in linkage disequilibrium (LD) with others, forming large haplotypes that pose challenges for editing and studying even in cultured human cells.

We reasoned that an approach considering the following points would be highly informative for determining the physiological roles and mechanisms of action for noncoding variants. First, despite limited sequence conservation, orthologous noncoding regions with similar local genomic organization across species likely influence biology through comparable mechanisms. Second, although the effect sizes of most noncoding variants are small in the general human population, they might be larger in subgroups of humans with susceptible genomic backgrounds and specific environmental exposures and, by extrapolation, in select animal models. Third, creating and analyzing isogenic human cells with precisely edited haplotypes, not just the sentinel SNPs, is crucial for definitively identifying effector genes for noncoding loci.

Based on these considerations, we have developed an efficient approach that integrates physiological studies in genetically engineered and phenotypically permissive animal models, precise editing of large haplotypes in human induced pluripotent stem cells (hiPSCs), and targeted chromatin conformation analysis. We utilized this approach to examine a 17.4 kbp noncoding genomic region containing 11 SNPs in linkage disequilibrium (LD) with the blood pressure (BP)-associated SNP rs1173771.^9^ High BP is the leading identifiable risk factor for disease burden and deaths worldwide.^10^ The study revealed robust effects of the orthologous noncoding region on BP in a rat model and identified the underlying physiological and molecular mechanisms. Our efficient approach can be applied to investigate any noncoding variants associated with human traits, providing new opportunities to overcome the crucial challenge of ascertaining the physiological roles and mechanisms of action for these variants.

## RESULTS

### An efficient, integrated, and targeted approach for investigating the physiological roles and mechanisms of action for long noncoding haplotype regions

Our approach to determining the physiological role and mechanisms of action for noncoding sequence variants begins by using whole genome sequence data to construct the haplotype in LD with a sentinel SNP identified by genome-wide association studies (GWAS) of a trait (**Fig. 1**). A region in the genome of model animals that is orthologous to the human noncoding haplotype region is identified through comparative mapping. This mapping takes into account sequence homology, the organization of neighboring genes, and any known epigenomic features. The orthologous noncoding region is then deleted in an animal strain with a genomic background susceptible to changes in the trait. These genetically engineered animals and their wild-type controls are subjected to relevant environmental stressors to provoke measurable differences in the trait, followed by examination of physiological and molecular mechanisms underlying these phenotypic differences (**Fig. 1**).

**Figure 1.**
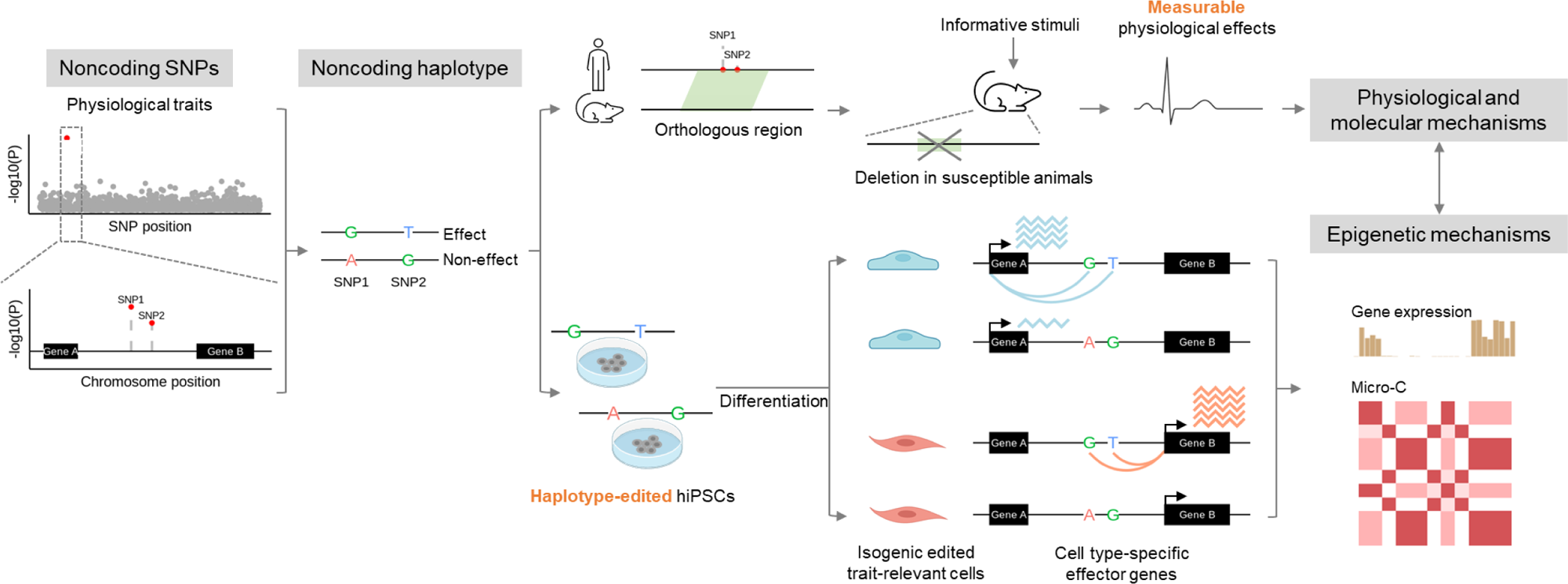
An efficient, integrated, and targeted approach to examining the physiological roles and mechanisms of action for noncoding haplotypes. SNP, single nucleotide polymorphism; hiPSCs, human induced pluripotent stem cells.

In parallel with physiological studies in animal models, we develop efficient genome-editing techniques to generate hiPSCs in which the entire reference and alternative haplotypes are reconstituted. These edited hiPSCs are differentiated into cell types relevant to the trait being studied. Differences in gene expression and epigenomic features are identified between isogenic cells containing either the reference or alternative haplotypes (**Fig. 1**). The findings from the human cells are cross-referenced with those from the animal models to interpret their physiological relevance.

### The rs1173771 locus associated with blood pressure

We applied the approach to study the rs1173771 locus. Several GWAS have identified robust associations of the SNP rs1173771 with human blood pressure.^9,11–13^ The G allele of rs1173771, which has a frequency of greater than 50% in several populations, is associated with higher mean arterial pressure (MAP), systolic and diastolic blood pressure (SBP, DBP), and pulse pressure (**Suppl. Table S1**).

We searched for SNPs that were in LD with rs1173771 using TOP-LD.^14^ TOP-LD is a tool for exploring LD inferred with high-coverage (∼30×) whole genome sequencing data from tens of thousands of individuals in the NHLBI Trans-Omics for Precision Medicine (TOPMed) program. We identified 10 SNPs in LD with rs1173771 at r^2^>0.8. The 11 SNPs span a 17.4-kbp region at nt 32,814,922 to 32,832,368 on human chr. 5 (hg38) (**Suppl. Table S2; Suppl. Fig. S1**).

The entire rs1173771 haplotype is in a noncoding, intergenic region of the human genome. The LD region is approximately 23 kbp downstream of the closest protein-coding gene *NPR3* (natriuretic peptide receptor 3) and 126 kbp from the *NPR3* transcription start site (**Suppl. Fig. S1**). The next closest protein-coding genes are *SUB1* (SUB1 regulator of transcription) and *TARS* (threonyl-tRNA synthetase 1), the transcription start sites of which are 283 kbp and 608 kbp from the rs1173771 LD region, respectively. Like most noncoding genomic loci associated with human traits, the physiological effect of the rs1173771 haplotype has not been examined with targeted manipulation of the genomic region.

### Identification and deletion of the orthologous genomic segment in Dahl salt-sensitive (SS) rats

The human *NPR3* and rodent *Npr3* loci and the surrounding genomic regions are highly syntenic. We identified a ∼30.4-kbp region in the rat genome (rn7 chr2: 60,825,801-60,856,216) that was orthologous to the human rs1173771 LD region (**Fig. 2A**; **Suppl. Fig. S2**). Similar to the human genome, the rat orthologous region was ∼14.5-kbp downstream of *Npr3* and ∼76-kbp from the annotated *Npr3* transcription start site, and *Sub1* and *Tars* are the next closest protein-coding genes.

**Figure 2.**
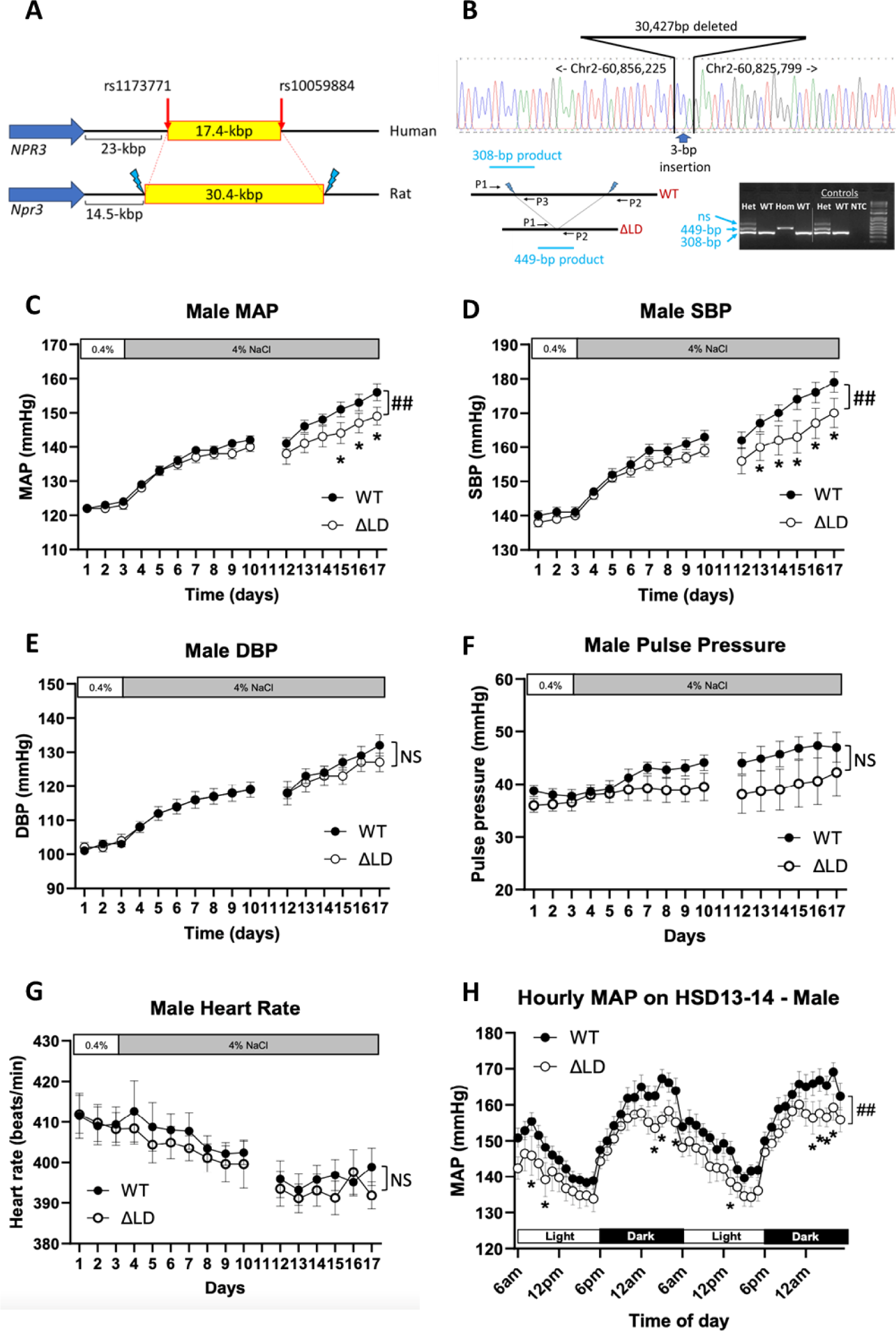
Deletion of the rs1173771 LD orthologous region attenuates salt-induced hypertension in male SS rats. **A.** Identification of the rat genomic region orthologous to the human rs1173771 LD (linkage disequilibrium) region. See Supplementary Figure S2 for more detail. **B.** Deletion of the rs1173771 LD orthologous region from the genome of SS rats. Daily averages of mean arterial pressure (MAP), systolic blood pressure (SBP), diastolic blood pressure (DBP), pulse pressure, and heart rate and hourly averages of MAP on days 13-14 of 4% NaCl high-salt diet (HSD) of male SS-Δrs1173771LD^−/−^ rats (ΔLD) and SS littermates (WT) are shown in panels **C, D, E, F, G,** and **H**. n=8; ##, p<0.01 for ΔLD vs. WT, two-way ANOVA repeated measures; *, p<0.05 for ΔLD vs. WT at the given time point, Holm-Sidak test.

We deleted the orthologous noncoding region in the Dahl salt-sensitive (SS) rat. The SS rat is a commonly used polygenic model of human salt-sensitive forms of hypertension.^15^ We reasoned that the genetic predisposition to the development of hypertension in SS rats and the high-salt diet as an established stressor for this model might amplify the effect of the noncoding genomic segment on blood pressure. We successfully deleted the 30.4-kbp orthologous region from the genome of the SS rat using CRISPR/Cas9 (**Fig. 2B**). The mutant strain was designated SS-Δrs1173771LD. The mutant rats bred without obvious health issues, and a 3-primer PCR could be used to distinguish zygosity at Mendelian ratios (**Fig. 2B**).

### Deletion of the rs1173771 LD orthologous region attenuates salt-induced hypertension in male SS rats

We investigated BP in SS-Δrs1173771LD^−/−^ and wild-type (WT) SS littermate rats (**Suppl. Fig. S3**). BP was not significantly different between SS-Δrs1173771LD^−/−^ rats and WT SS littermates on a 0.4% NaCl diet. BP was increased in both SS-Δrs1173771LD^−/−^ rats and SS littermates after the rats were switched to a 4% NaCl high-salt diet (HSD). However, the 24-hour MAP was significantly lower in male SS-Δrs1173771LD^−/−^ rats than in male SS rats after 12 days on the HSD (**Fig. 2C**). After 14 days on the HSD, MAP was 149.1 ± 2.7 mmHg in SS-Δrs1173771LD^−/−^ rats, compared with 155.9 ± 2.5 mmHg in SS rats (n=8, p<0.05). The difference in MAP could be attributed to the difference in SBP (**Fig. 2D**). SBP was 169.6 ± 4.3 mmHg in SS-Δrs1173771LD^−/−^ rats after 14 days on the HSD, compared with 179.0 ± 3.0 mmHg in littermate SS rats (n=8, p<0.05). DBP, pulse pressure, and heart rate were not significantly different between SS-Δrs1173771LD^−/−^ and SS rats (**Fig. 2E-2G**). The difference in MAP was observed throughout the day and was particularly prominent in the later stage of the dark (active) period (**Fig. 2H**).

BP of female rats were measured as well. No difference was observed between female SS-Δrs1173771LD^−/−^ and SS (**Suppl. Fig. S4**).

### Deletion of the rs1173771 LD orthologous region leads to upregulation of arterial Npr3 expression and improves CNP-induced vasodilation

RNA-seq was performed to identify any genes that might be influenced by the deletion of the rs1173771 LD orthologous region. In mesentery artery, Npr3 expression was elevated by about 50% in male SS-Δrs1173771LD^−/−^ rats while expression of neighboring genes Sub1 and Tars remained unchanged (**Fig. 3A**). *Npr3* encodes natriuretic peptide receptor 3, also called atrial natriuretic peptide receptor C (NPR-C). Western blot analysis confirmed the upregulation of NPR-C protein level in the mesentery artery in male SS-Δrs1173771LD^−/−^ rats (**Fig. 3B**). Immunofluorescence analysis further confirmed the upregulation of NPR-C and showed that the upregulation occurred in both the endothelia and the smooth muscles (**Fig. 3C**).

**Figure 3.**
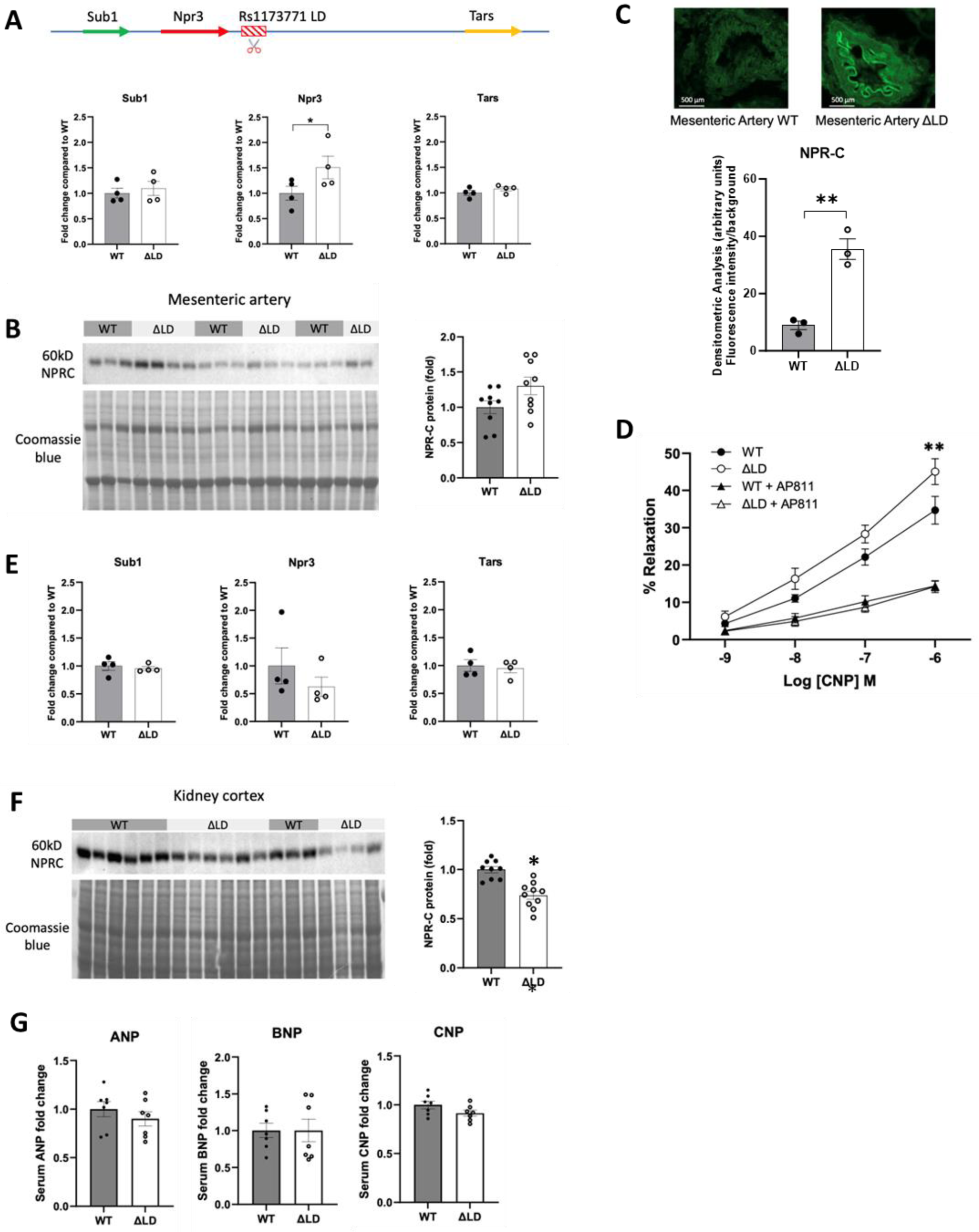
Deletion of the rs1173771 LD orthologous region leads to upregulation of arterial Npr3 expression and improves CNP-induced vasodilation. **A.** Upregulation of Npr3 in mesentery arteries in male SS-Δrs1173771LD^−/−^ rats (ΔLD) compared with SS littermates (WT), based on RNA-seq. The schematic at the top shows the genomic positions of the rs1173771 LD orthologous region relative to Npr3, Sub1, and Tars genes. N=4; *, p<0.05, unpaired t-test. **B.** Western blot analysis of NPR-C in mesentery arteries from male ΔLD and WT rats. N=9; p=0.06, unpaired t-test. **C.** Immunofluorescence analysis of NPR-C in mesentery arteries from male ΔLD and WT rats. N=3; **, p<0.01, unpaired t-test. **D.** Vasodilation response of mesentery arteries from male ΔLD and WT rats in response to increasing concentrations of C-type natriuretic peptide (CNP). AP811, an NPR-C antagonist, was used at 100nM. N=6; **, p<0.01 for ΔLD vs. WT by Holm-Sidak test. **E.** Npr3 expression in the kidney cortex in male ΔLD and WT rats based on RNA-seq. N=4. **F.** Western blot analysis of NPR-C in the kidney cortex from male ΔLD and WT rats. N=9 WT and 10 ΔLD; *, p<0.05, unpaired t-test. **G.** Serum levels of atrial, B-type, and C-type natriuretic peptides (ANP, BNP, CNP). N=7.

NPR-C has multiple functions. In the vasculature, NPR-C binds C-type natriuretic peptide (CNP), mediates the effect of CNP on vascular function and structure, and plays an important role in preserving vascular homeostasis in vivo.^16,17^ We found that vasodilation of mesenteric arteries in response to CNP was significantly enhanced in male SS-Δrs1173771LD^−/−^ rats compared with SS littermates (**Fig. 3D**). AP811, a selective NPR-C antagonist, attenuated vasodilation in both groups and abolished the difference of vasodilation between the two groups, suggesting that the enhancement of CNP-induced vasorelaxation in vessels from SS-Δrs1173771LD^−/−^ rats was NPR-C dependent (**Fig. 3D**).

In the kidney, NPR-C facilitates the clearance of atrial, B-type, and C-type natriuretic peptides (ANP, BNP, CNP) from the circulation via endocytosis.^18,19^ Natriuretic peptides have broad and potent physiological effects including promoting natriuresis and lowering blood pressure. RNA-seq analysis of the kidney cortex indicated that Npr3 expression tended to be down-regulated in male SS-Δrs1173771LD^−/−^ rats (**Fig. 3E**). Western blot analysis showed significant down-regulation of NPR-C in the kidney cortex (**Fig. 3F**). However, serum levels of ANP, BNP, and CNP were not significantly different between male SS-Δrs1173771LD^−/−^ rats and SS littermates (**Fig. 3G**).

Female rats showed a similar trend as male rats in that Npr3 expression in SS-Δrs1173771LD^−/−^ rats tended to be higher in mesenteric arteries and lower in the kidney cortex compared with the wild-type SS littermates, although the differences did not reach statistical significance (**Suppl. Fig. S5**).

### Deletion of the rs1173771 haplotype region influences gene expression in iECs and iVSMCs

Next, we examined the effect of the rs1173771 haplotype on gene expression in human cell types relevant to BP regulation. Urine cells obtained from two individuals were successfully reprogrammed to generate iPSC lines Y4 and 39b. Both lines had robust expression of markers of pluripotent stem cells and exhibited the expected morphology and normal karyotype (**Suppl. Fig. S6A-S6D**). The source subjects for Y4 and 39b had hypertension and normotension, respectively, at the time of urine collection. DNA sequencing showed that Y4 had homozygous BP-elevating alleles at all 11 SNPs in the rs1173771 haplotype, and 39b was heterozygous for the 11 SNPs (**Suppl. Fig. S6E, S6F; Suppl. Table S2**).

The large, 17.4-kbp size of the rs1173771 haplotype region and the presence of 11 SNPs in LD make it particularly challenging to edit this haplotype. We tested several approaches and found a two-step, CRISPR-Cas9 mediated genome editing approach to be the most efficient for reconstituting homozygous BP-elevating or -lowering rs1173771 haplotype in hiPSCs (**Fig. 4**). The first step of this approach was to delete the entire haplotype region. Two gRNAs flanking the rs1173771 haplotype region were used to delete the 17.4-kbp region in the iPSC lines. Screening of 52 clones of Y4 cells and 103 clones of 39b cells following transfection identified 3 and 2 clones, respectively, with successful homozygous deletion of the haplotype region. The deletion was confirmed by PCR and sanger sequencing (**Fig. 5A-5E**), as well as whole genome sequencing. These cell clones were designated Y4 ΔLD and 39b ΔLD. Y4 ΔLD and 39b ΔLD cells retained robust expression of markers of pluripotency and exhibited the expected morphology and normal karyotype (**Suppl. Fig. S7**).

**Figure 4:**
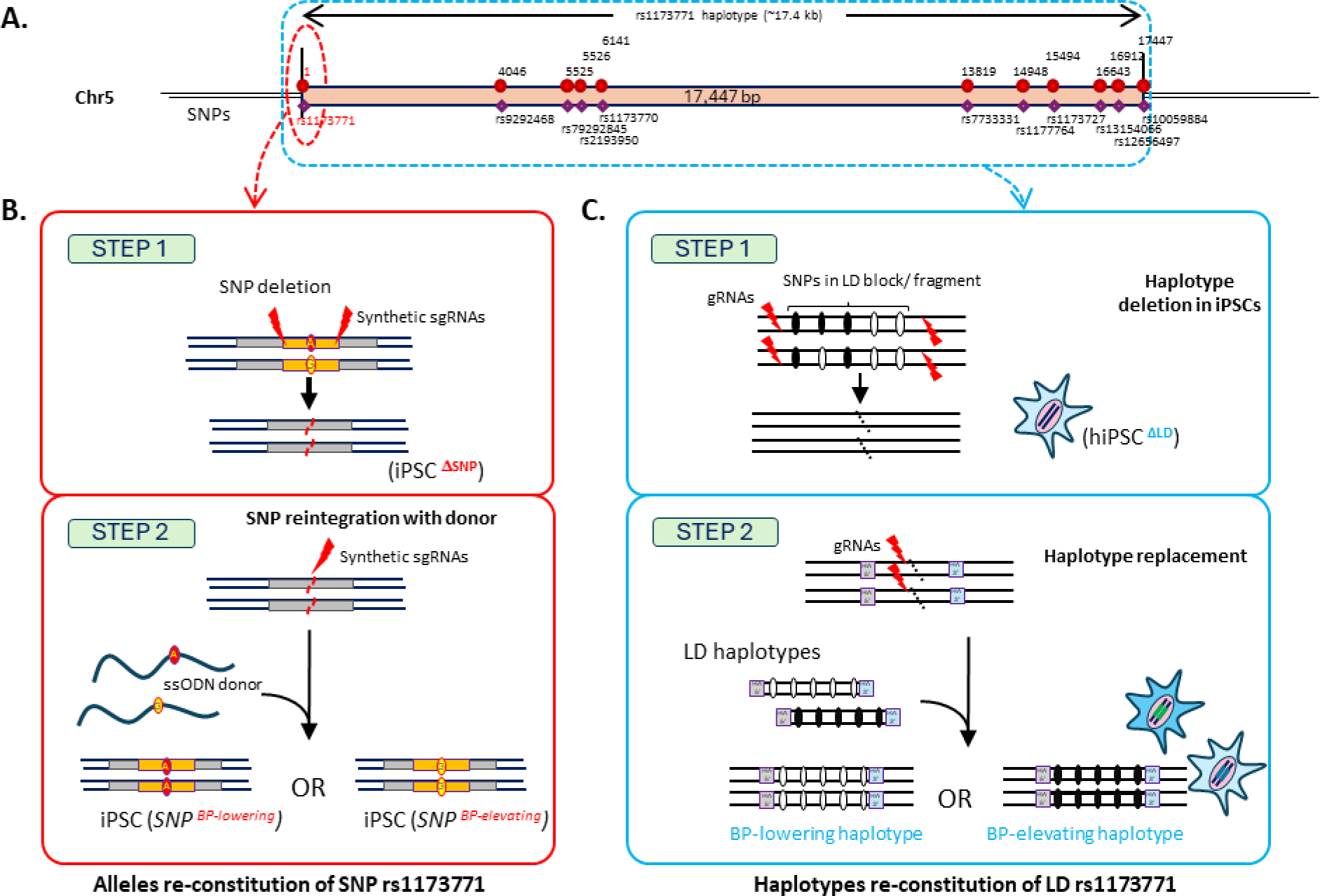
An efficient, two-step genome editing approach for generating isogenic iPSCs with homozygous alleles or haplotypes. **A.** The rs1173771 haplotype contains 11 SNPs spanning 17.4 kbp. Deletion and re-constitution approaches for generating isogenic human iPSC lines with homozygous alleles at the single SNP rs1173771 **(B)** and all 11 SNPs in the rs1173771 haplotype **(C)** are shown.

**Figure 5:**
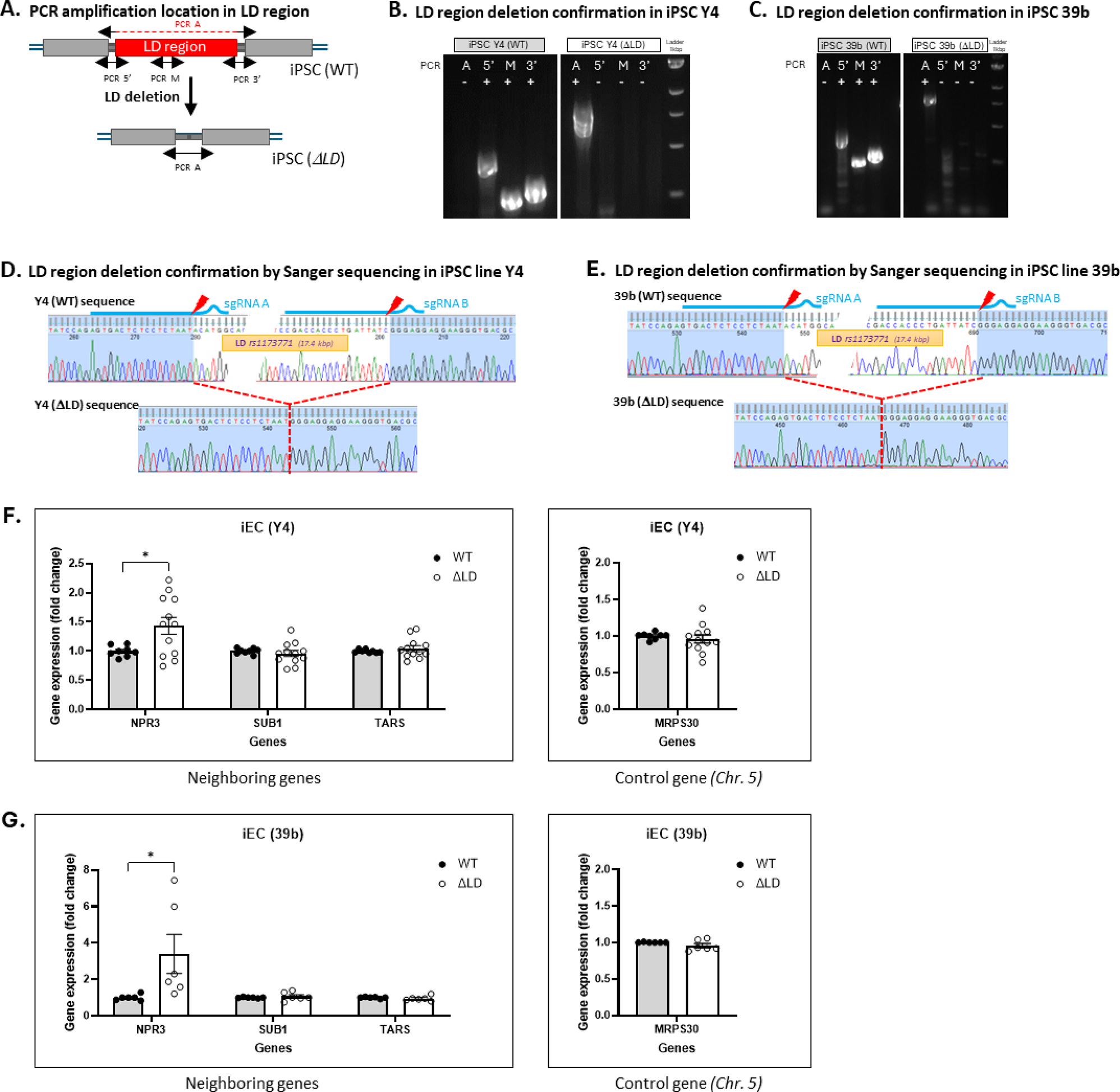
Deletion of the rs1173771 haplotype region results in upregulation of NPR3 in iECs and iVSMCs. **A.** Locations of various PCR amplicons relative to the rs1173771 haplotype region. **B-E.** Confirmation of rs1173771 haplotype region deletion in Y4 and 39b cells by PCR (B, C) and Sanger sequencing (D, E). **F.** mRNA expression of neighboring genes NPR3, SUB1, and TARS in endothelial cells (CD144^+^) differentiated from iPSC Y4 (WT) and rs1173771 LD region deleted Y4 (ΔLD). n=8 (WT) and n=12 (ΔLD) independently differentiated samples from 3 rounds of differentiations. *p ≤ 0.05, unpaired t-test. **G.** mRNA expression of NPR3, SUB1, and TARS in endothelial cells (CD144^+^) differentiated from iPSC 39b (WT) and rs1173771 LD region deleted 39b (ΔLD). n=6 (WT) and n=6 (ΔLD) independent differentiated samples from 3 rounds of differentiations. *p ≤ 0.05, unpaired t-test.

Y4 ΔLD and 39b ΔLD cells, along with wild-type Y4 and 39b cells, were differentiated to endothelial cells (iEC). The iECs showed robust expression of EC marker genes (**Suppl. Fig. S8**). Compared with iEC derived from wild-type Y4 and 39b cells, iECs derived from Y4 ΔLD and 39b ΔLD cells showed significantly higher abundance of NPR3 mRNA expression (**Fig. 5F, 5G**). The abundance of neighboring genes SUB1 and TARS, as well as a distant gene MRPS30 (∼12 Mbp from the rs1173771 haplotype region), was not affected by the deletion. These results were consistent with the findings in mesentery arteries in SS-Δrs1173771LD^−/−^ rats described above.

### Effect of BP-elevating and -lowering alleles of the rs1173771 haplotype on gene expression in isogenic iECs and iVSMCs

One clone of Y4 ΔLD and 39b ΔLD cells were subject to Step 2 editing to generate iPSCs with homozygous BP-elevating or -lowering alleles for all 11 SNPs in the rs1173771 haplotype region (**Fig. 4**). This involved the use of 19.6 kbp long constructs containing homozygous BP-elevating or -lowering alleles at all 11 SNPs as donor constructs for homology-dependent repair editing. Screening of 611 clones from Y4 ΔLD following transfection identified 1 and 6 clones containing homozygous BP-elevating and BP-lowering alleles, respectively, at all 11 SNPs (**Fig. 6**). One clone with each genotype was designated ‘Y4 BP-elevating haplotype’ and ‘Y4 BP-lowering haplotype’ and used in subsequent studies. Screening of 567 clones from 39b ΔLD following transfection identified 1 and 1 clones containing homozygous BP-elevating and BP-lowering alleles, respectively, at all 11 SNPs (**Fig. 6**). These clones were designated ‘39b BP-elevating haplotype’ and ‘39b BP-lowering haplotype’ and used in subsequent studies. These edited iPSCs retained robust expression of stem cell markers and showed expected morphology and normal karyotype (**Suppl. Fig. S9**).

**Figure 6:**
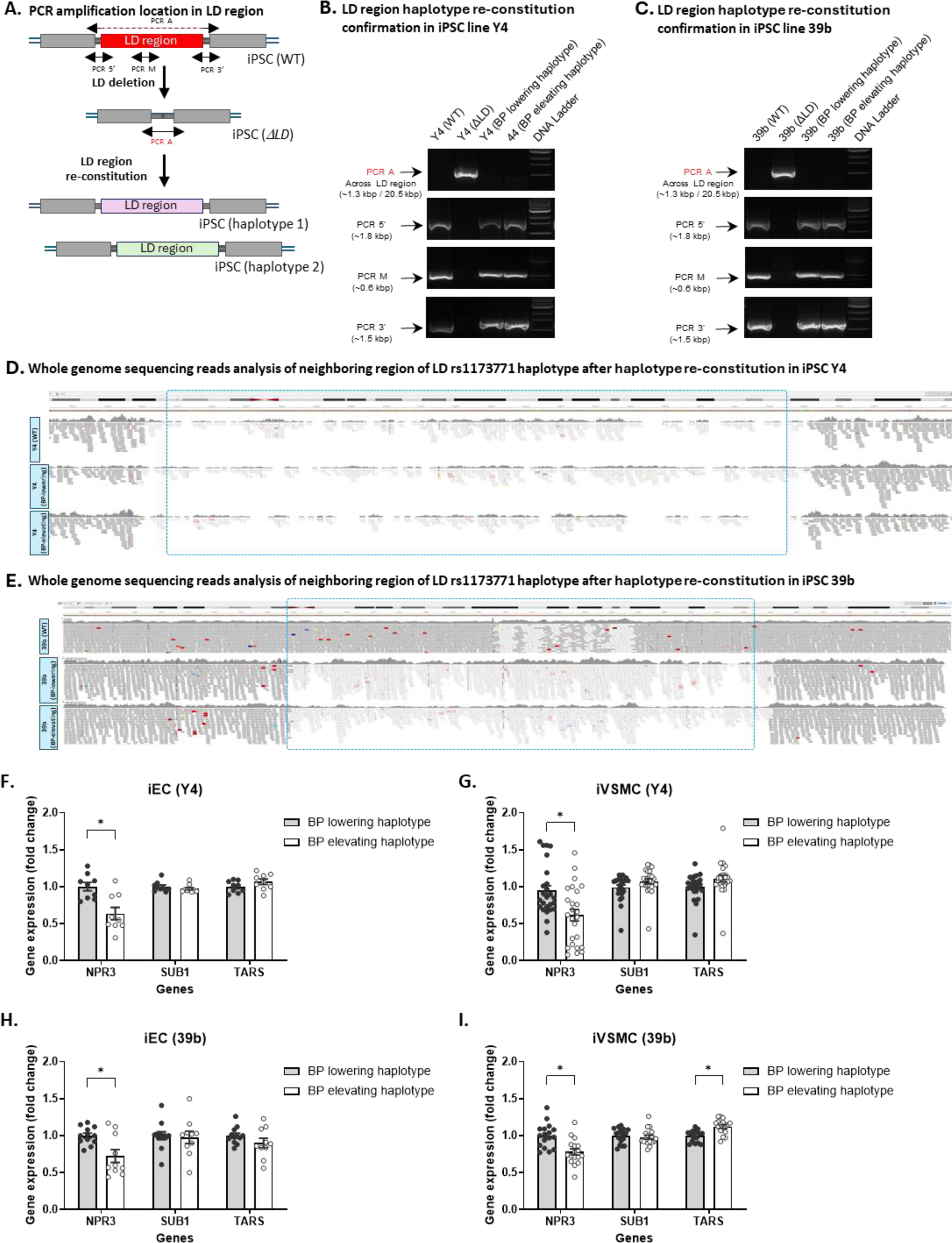
The BP-lowering rs1173771 haplotype increases NPR3 expression in iECs and iVSMCs. **A.** Locations of various PCR amplicons relative to the rs1173771 haplotype region. Confirmation of rs1173771 haplotype region re-insertion in Y4 (**B**) and 39b (**C**). Whole genome sequencing reads at the rs1173771 haplotype and surrounding regions showing the reconstitution of homozygous BP-elevating or -lowering haplotype [Chr5 pos (hg38) 32,814,922-32,832,368] in iPSC lines Y4 **(D)** and 39b **(E).** Expression of neighboring genes NPR3, SUB1, and TARS in endothelial cells (iEC) and vascular smooth muscle cells (iVSMC) differentiated from iPSCs containing BP-lowering and -elevating haplotypes in Y4 (**F, G**), and 39b (**H, I**). N=9-25 independently differentiated samples from 3-5 rounds of differentiation. *p ≤ 0.05, unpaired t-test.

We differentiated Y4 BP-elevating haplotype, Y4 BP-lowering haplotype, 39b BP-elevating haplotype, and 39b BP-lowering haplotype to iECs and iVSMCs. The iECs and iVSMCs showed robust expression of their respective marker genes (**Suppl. Fig. S10**). Compared to iECs and iVSMCs with the BP-lowering rs1173771 haplotype, iECs and iVSMCs with the BP-elevating rs1173771 haplotype showed significantly lower levels of NPR3 expression (**Fig. 6F-6I**). The expression of neighboring genes SUB1 and TARS was not different between iECs with BP-lowering or -elevating haplotypes. In iVSMCs, SUB1 expression was not different between haplotypes, while TARS showed a slight, 10% higher expression with the BP-elevating haplotype (**Fig. 6F-6I**).

### Effect of BP-elevating and -lowering alleles of the single rs1173771 SNP on gene expression in isogenic iECs and iVSMCs

In a separate series of two-step genome editing experiments (**Fig. 4**), we generated 39b cells with homozygous BP-lowering or -elevating alleles of the single SNP rs1173771, which was the sentinel SNP reported in GWAS of BP. Successful editing was confirmed by sequencing (**Fig. 7A-7E**).

**Figure 7:**
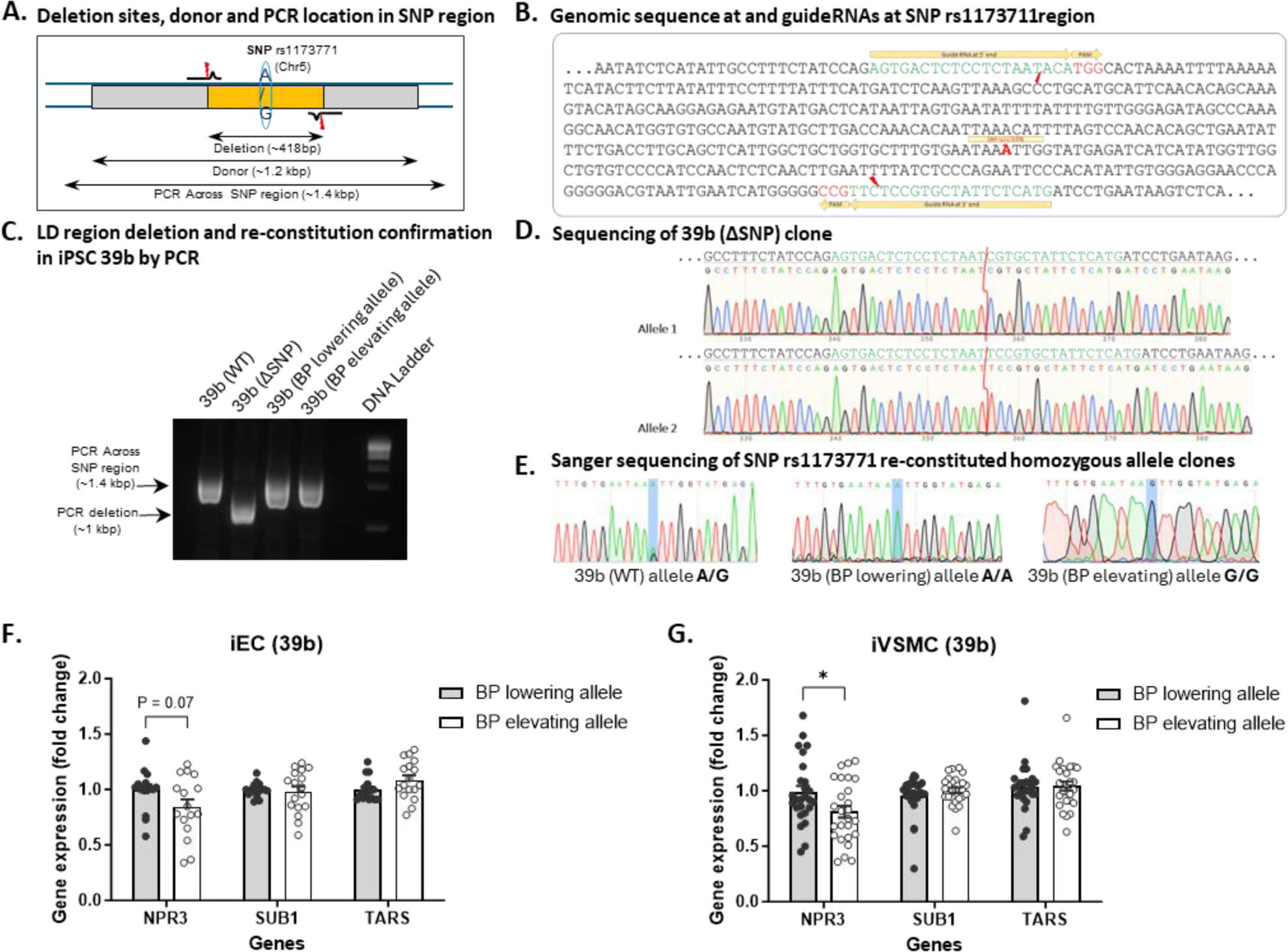
Effect of the single SNP rs1173771 on the expression of neighboring genes. **A.** Locations of various PCR amplicons relative to the rs1173771 single SNP region. **B.** gRNAs locations. Confirmation of rs1173771 single SNP region deletion and reconstitution of homozygous BP-elevating and –lowering alleles in 39b cells (**C, D, E**). Expression of neighboring genes NPR3, SUB1, and TARS in endothelial cells (iEC) (**F**) and vascular smooth muscle cells (iVSMC) (**G**) differentiated from iPSCs containing homozygous BP-lowering and -elevating alleles at the single SNP rs1173771. n=17-26 independently differentiated samples from 4-5 rounds of differentiation. *p ≤ 0.05, unpaired t-test.

In iECs derived from these edited cells, NPR3 levels tended to be lower in cells with homozygous BP-elevating allele at the single SNP rs1173771 compared with cells with homozygous BP-lowering allele, but the difference did not reach statistical significance (**Fig. 7F**). The average reduction of NPR3 expression in iECs with homozygous BP-elevating allele at rs1173771 was 16%, compared with 27% in iECs with homozygous BP-elevating alleles for the entire rs1173771 haplotype including 11 SNPs (see Fig. 6H). In iVSMCs, NPR3 was significantly down-regulated in cells with homozygous BP-elevating allele at rs1173771 (**Fig. 7G**). SUB1 and TARS were not differentially expressed in either iECs or iVSMCs (**Fig. 7F, 7G**).

### Effect of BP-elevating and -lowering alleles of the rs1173771 haplotype on chromatin interactions in isogenic iECs and iVSMCs

The rs1173771 haplotype region is 126 kbp from the NPR3 transcription start site, suggesting that the effect of the haplotype on NPR3 expression is likely mediated by chromatin folding and interaction. We hypothesized that the BP-lowering and BP-elevating haplotypes of rs1173771 would have differential effects on the chromatin interaction between the haplotype region and the *NPR3* promoter. We tested this hypothesis using region capture Micro-C with probes designed to cover the rs1173771 haplotype region containing SNPs and the promoter regions of neighboring genes including *SUB1*, *NPR3*, and *TARS* (**Suppl. Fig. S11**). The assay was performed in iECs and iVSMCs containing homozygous BP-lowering or -elevating rs1173771 haplotype. The assay generated 4.6 to 8.1 million non-duplicated read pairs in each sample, approximately 90% of which were valid read pairs (**Suppl. Table S3**).

In both iECs and iVSMCs, the BP-elevating rs1173771 haplotype increased chromatin contacts between the rs1173771 haplotype region and *NPR3* promoter region, compared to the BP-lowering haplotype (**Fig. 8A-8D**). Chromatin contacts between the haplotype region and *SUB1* or *TARS* promoter region were similar between cells containing the BP-lowering and -elevating haplotypes.

**Figure 8:**
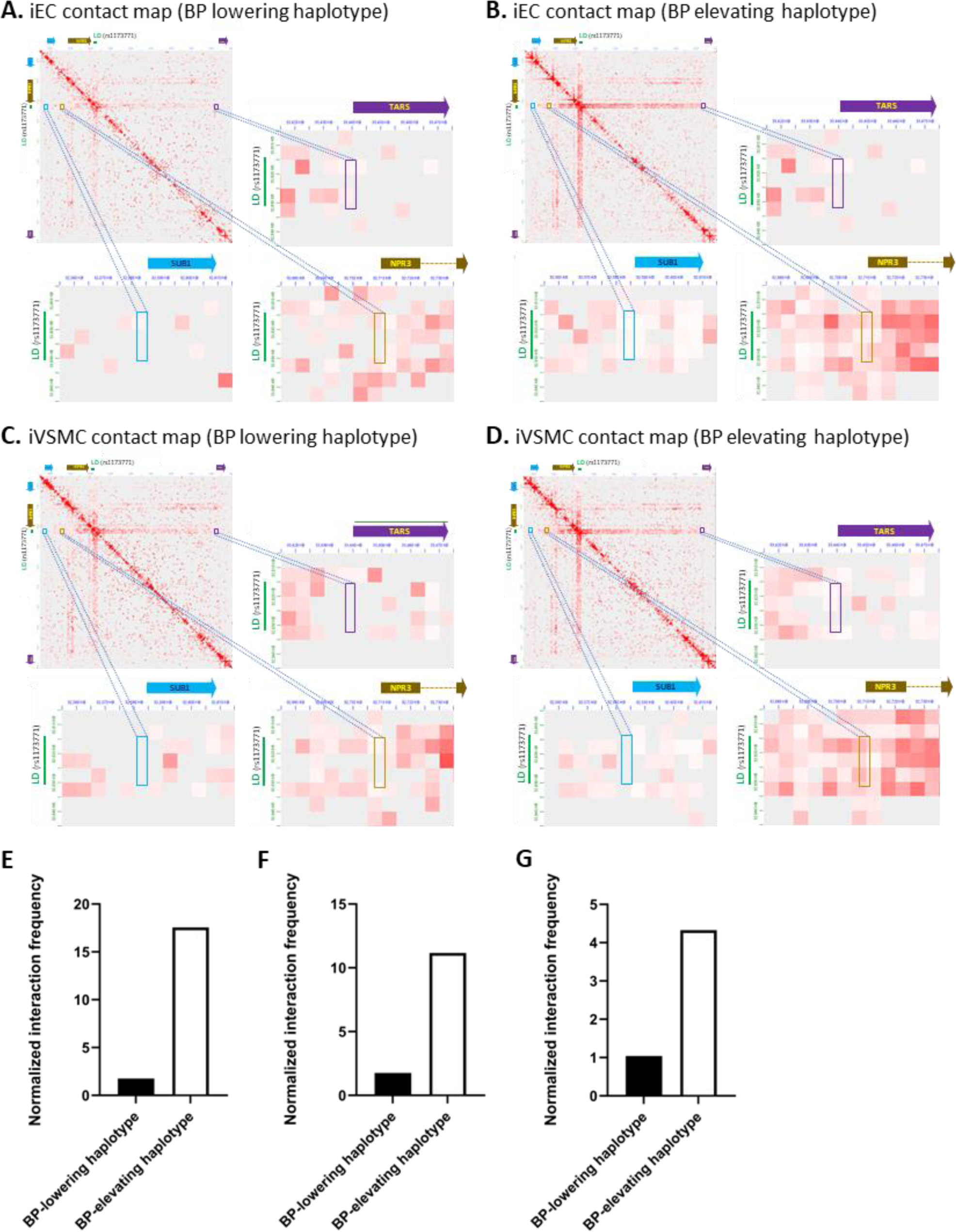
The BP-elevating rs1173771 haplotype interacts more frequently with the NPR3 promoter region than the BP-lowering rs1173771 haplotype. Chromatin contact maps at rs1173771 and the surrounding region, based on a region capture Micro-C analysis, are shown for iECs with BP-lowering rs1173771 haplotype (**A**), iECs with BP-elevating rs1173771 haplotype (**B**), iVSMCs with BP-lowering rs1173771 haplotype (**C**), and iECs with BP-elevating rs1173771 haplotype (**D**). The zoom-in images in each panel show chromatin contacts between the rs1173771 haplotype region with promoters of SUB1, NPR3, and TARS. Note the greater contacts for the NPR3 promoter shown in panel B vs A and D vs. C. Significantly different, normalized chromatin contact frequencies are shown for regions overlapping the rs1173771 haplotype with the NPR3 promoter region in iECs (**E**; p.adj = 0.009) and a region upstream of the NPR3 promoter in iECs (**F**; p.adj = 0.001) and iVSMCs (**G**; p.adj = 0.02). The sequence coordinates for these regions are available in Supplementary Table S4. The statistical significance was based on HiCCompare analysis with adjusted p < 0.05.

We performed a formal statistical comparison of chromatin interaction using HiCCompare. In iECs, the interactions of a genomic segment overlapping the 5’ end of the rs1173771 haplotype region with the *NPR3* promoter region and a region just upstream of the promoter were significantly stronger in iECs with homozygous BP-elevating haplotype compared to BP-lowering haplotype (**Fig. 8E, 8F; Suppl. Table S4**). Similarly, in iVSMCs, the interaction of a genomic segment overlapping the rs1173771 haplotype region with a region just upstream of the *NPR3* promoter was significantly stronger for homozygous BP-elevating haplotype compared to BP-lowering haplotype (**Fig. 8G; Suppl. Table S4**). No significant difference was found for the interactions between the haplotype region and *SUB1* or *TARS* promoter region in either iECs or iVSMCs.

These results suggest that the BP-elevating rs1173771 haplotype interacts with the *NPR3* promoter more strongly to suppress NPR3 expression, possibly through the recruitment of inhibitory protein complexes. NPR3 expression is up-regulated when the interaction of *NPR3* promoter with the haplotype region is weakened by the BP-lowering haplotype or lost when the haplotype region is deleted.

## DISCUSSION

This study has shown the feasibility of using genome editing in animal models to demonstrate the physiological role of a noncoding genomic region associated with human traits. The deletion of the rs1173771 LD orthologous region resulted in a 9.4 mmHg reduction in SBP in SS rats on a 4% NaCl diet for 14 days. This reduction in SBP is remarkably large as the effect size of the rs1173771 locus on the SBP of the general human population was only approximately 0.5 mmHg.^11^ Our study was performed in a rat strain with a genomic background that predisposes the rat to the development of salt-sensitive hypertension.^15^ This genomic background and the use of a high-salt diet as a “second hit” may have amplified the effect of deleting the rs1173771 LD orthologous region on BP. The findings suggest that in individuals with specific genomic backgrounds and exposed to certain environment and lifestyle factors, the rs1173771 locus may have greater effects on BP than what the GWAS suggested.

The rs1173771 LD orthologous region deletion influenced BP only in male rats. In global Npr3 gene knockout mice, BP was increased in female mice and decreased in male mice, and impaired vasodilation was observed only in female mice.^16^ In the current study, deleting the rs1173771 LD orthologous region resulted in changes in Npr3 expression similarly in male and female rats. Therefore, the male-specific effect of rs1173771 LD orthologous region deletion on BP may not be due to any sex-specific effect of the noncoding genomic region on Npr3 expression but rather due to sex-specific effects of the NPR-C mechanism, which warrant further studies.

The rs1173771 LD orthologous region deletion increased NPR3 expression in mesentery arteries and decreased it in the kidney cortex. The tissue-specific effect is consistent with recent studies that have identified the BP-associated rs1173771-G allele as an eQTL for driving increased^20^ or decreased^21^ NPR3 expression in different tissues. Additional studies are needed to investigate the mechanisms underlying the tissue-specific effect.

Our hiPSC study is one of the first to specifically edit a large noncoding haplotype and identified the effect of the haplotype on target gene expression in cell types relevant to the associated trait. The two-step genome editing strategy is sufficiently efficient to enable the precise editing of 11 SNPs spanning a 17.4 kbp haplotype region. Several clones of hiPSC with homozygous deletion of the region and precise reconstitution of homozygous haplotypes were obtained from the screening of a few dozen and a few hundred clones, respectively. Rs1173771 and a few other SNPs in the haplotype have been identified as eQTLs for *NPR3* in several tissues.^5,22^ In an elegant study by Ren, et al, some BP-elevating alleles in the rs1173771 LD block were associated with lower NPR3 expression in primary vascular smooth muscle cells (VSMCs) obtained from 100 individuals.^21^ Our large haplotype editing approach enables direct comparisons of isogenic cells and provides definitive proof that the BP-elevating rs1173771 haplotype sequence, compared with the BP-lowering haplotype sequence, decreases NPR3 expression in iECs and iVSMCs. These findings, together with the findings from iECs with the rs1173771 haplotype region deleted, reinforces the findings from the animal models and supports the notion that the rs1173771 haplotype region influences NPR3 expression and, thereby, affects blood pressure.

The allelic effect of the single SNP rs1173771 does not fully recapitulate the allelic effect of the rs1173771 haplotype that includes 11 SNPs in iECs, although it appears to do so in iVSMCs. These findings indicate that the biological effect of a SNP may be influenced by the presence of other SNPs in LD with it. Moreover, the relative contributions of the various SNPs in a haplotype to a biological effect may vary depending on the cell type.

We are cautious in extrapolating the effect of the rs1173771 LD orthologous region deletion in rats to the effect of the specific SNPs in the rs1173771 haplotype in humans. It is challenging to pinpoint the specific nucleotides in the rat genome that correspond to the SNPs in the rs1173771 haplotype because the noncoding genomic region is not fully conserved between rat and human. It would be interesting to reconstitute the human haplotype in the rat genome and examine its physiological effects, but it is challenging to do so because of the large size of the haplotype. Nevertheless, it is plausible that the rs1173771 LD orthologous region in rats and the human rs1173771 haplotype region regulate effector genes through similar epigenetic mechanisms including chromatin interaction. The findings of the current study support an important role for these epigenetic mechanisms in regulating NPR3 expression, vascular function, and blood pressure.

The feasibility of applying the approach established in this study to examine other noncoding variants and haplotypes is high. Orthologous regions in rodent genomes can be identified for a substantial portion of human noncoding variants.^23^ The orthologous regions can be deleted efficiently for in vivo investigation, and our two-step genome editing method is efficient for reconstituting even large haplotypes in cultured cells. Application of this efficient, integrated, and targeted approach should contribute significantly to closing the critical gap between genetic discoveries and physiological and pathophysiological understanding.

## METHODS

### Genomic segment deletion in SS rats

CRISPR/Cas9 methods were used to delete a noncoding genomic segment in the SS (SS/JrHsdMcwi) rat. The LiftOver tool at UCSC Genome Browser placed the non-coding syntenic region to the human haplotype (defined in **Suppl. Table S2 and Fig. S1**) at mRatBN7.2 chr2:60,826,449-60,855,714, a 30.4-kbp segment of chromosome 2 distal to the rat *Npr3* gene. spCas9 (QB3 MacroLab, University of California, Berkely) was mixed with dual pairs of sgRNAs (Integrated DNA Technologies) flanking this region, targeting the sequences ATGAAGTGCCTAGCTTCATT, GAAGCTAGGCACTTCATCTA, ATTCATAATGGGACACTCGA, and CCCAGCCTCAGATTCATAAT and injected into fertilized SS/JrHsdMcwi strain rat embryos. A founder was identified harboring a 30.4-kbp deletion (mRatBN7.2 chr2:60,856,216-60,825,801) along with TGG insertion and was backcrossed to the parental strain to establish a breeding colonyu. This SS-Δrs1173771LD^−/−^ strain is registered as SS-Del(2q)1Mcwi (RGDID: 401938598). Heterozygous SS-Δrs1173771LD^+/−^ breeders were maintained on a purified 0.4% NaCl diet AIN-76A (Dyets, Inc).

### Blood pressure measurement using radiotelemetry

Continuous measurement of blood pressure in conscious, freely moving rats was performed using radiotelemetry as described (**Suppl. Fig. S3**). ^24,25^ Briefly, a telemetry transmitter (model HD-S10, Data Systems International) was implanted in the right carotid artery when rats were 7 weeks of age. After one week of recovery, continuous BP monitoring was started. Following three days of stable baseline BP recording, the diet was switched to an AIN-76A diet containing 4% NaCl (high salt diet, HSD) for 14 days. Tissues and serum were collected at the end of the experiment. Both male and female rats were studied.

### RNA-seq

RNA-seq experiment and data analysis were performed as described.^26,27^

### Western blot

Western blotting was performed as descried.^25,28^ Primary antibody for rat NPRC was obtained from Novus Biologicals (catalog number NBP1-90163) and used at 1:1000 dilution. Secondary antibody was Anti-Rabbit IgG (Sigma A6154) and used at 1:5000.

### Immunofluorescence staining in mesentery artery

Optimal cutting temperature compound (OCT) embedded fresh tissue samples were subject to frozen sectioning and 5 µm thick sections were mounted onto slides for immunohistochemistry. Slides were brought to room temperature and then fixed in a 1:1 solution of methanol/acetone (v/v) for 10 minutes at −20C. The tissues were washed twice with 1X PBS for 2 minutes each. The tissues were permeabilized in a solution of 0.1% trition X 100 in 1XPBS for 10 minutes at room temperature. After 1 wash with 1XPBS the tissues were blocked in 2.5% horse serum for 20 minutes at room temperature. Next, the tissues were incubated with anti-NPRC antibody^29^ at a dilution of 1:250 for 30 minutes at room temperature. The tissues were washed 3 times in 1XPBS for 2 minutes each. The tissues were incubated with secondary antibody (Vector Labs) for 30 minutes at room temperature. After 3 washes with 1X PBS the tissues were cover slipped using anti-fade mounting media (Vector Labs). The slides were imaged on an Olympus BX41 microscope using a 40X objective. Image J (NIH) was used to quantify the fluorescence intensity.

### Vasoreactivity analysis

Vasoreactivity analysis was performed as described.^26,30^ Briefly, third order mesenteric artery from naïve male rats were used in the current study. Isolated artery segments were mounted between two glass cannulas in a culture myograph chamber (DMT, 204CM). After equilibration, U46619 (Catalog No. 16450, Cayman Chemical) was used to induce vessel contraction. Once a stable response to U46619 was achieved, 0.001-1μM CNP (Catalog No. N8768, Sigma-Aldrich) was administrated to construct cumulative concentration-response curves in the absence or presence of 100nM AP 811 (Catalog No. 5498, TOCRIS), a selective NPR-C antagonist.

### Natriuretic peptide assay

Analysis of serum levels of ANP, BNP, and CNP was performed using ELISA assays (EIA-ANP and EIAM-BNP from RayBiotech, CNP-22 EIA kit EKE-012-03 from Phoenixpeptide) following the manufacturer’s instructions.

### iPSCs and genome editing of long haplotype or single SNP

Two iPSC lines were reprogrammed from urine cells obtained from well-phenotyped individuals as we described.^31,32^ A two-step genome editing strategy was designed and applied to these iPSC lines to generate isogenic iPSCs with homozygous BP-elevating or BP-lowering alleles for the single SNP rs1173771 or the 11 SNPs in the rs1173771 haplotype region that spans 17.4 kbp (**Fig. 1**).

For the first step of 2-step single SNP editing, two synthetic sgRNAs targeting the 20-200 bp region at the 3’- and 5’-end of the SNP rs1173771 (**Suppl. Table S5**) and spCas9-2NLS Nuclease protein were used to delete a segment of DNA around rs1173771. After 48 hrs of transfection, cells were dissociated for single cell cloning using Microfluidic-based benchtop single cell sorter HANA (Namocell). Single cell clones were further grown for 10-14 days and characterized for the deletion of SNP region using quick extraction of DNA and PCR (**Suppl. Table S6**). iPSC clones showing the deletion of SNP region with shorter PCR fragment were further characterized for deletion of the region around rs1173771 by PCR, sanger sequencing and whole genome sequencing. iPSC line with the deletion of the region around rs1173771 was tested for their pluripotency status, differentiation potential and genetic stability by molecular karyotyping. For the second step of 2-step SNP editing, homology dependent knock-in approach was used which required a gRNA plasmid targeting the junction of single SNP rs1173771 region deletion site (**Suppl. Table S5**) and ssDNA donor fragment for BP-elevating rs1173771-G or BP-lowering rs1173771-A alleles with approximately 500-600 bp long homology arm at the 3’- and 5’-end of the SNP rs1173771 (**Suppl. Table S6**). Isogenic iPSC lines with the insertion of homozygous BP-elevating rs1173771 and BP-lowering rs1173771 alleles were selected, confirmed, and assessed for quality as described above.

A modified 2-step approach was used for long haplotype editing. In the first step, two gRNA plasmids targeting the 3’- and 5’-end of the haplotype region (**Suppl. Table S7**) were used for the deletion of the 17.4 kbp rs1173771 haplotype region. Single cell clones with successful deletion of the haplotype region were selected, confirmed, and assessed for quality as described above (**Suppl. Table S6**). For the second step of 2-step haplotype editing, homology dependent knock-in approach was used which required a gRNA plasmid targeting at the junction of haplotype region deletion site (**Suppl. Table S7**) and a donor plasmid for BP-elevating rs1173771 haplotype region or BP-lowering rs1173771 haplotype region with approximately 1 kb long homology arm at the 3’- and 5’-end of the haplotype region. For the donor plasmid preparation, the rs1173771 haplotype region with homology arm region (total of ∼20 kb long) was amplified from DNA isolated from iPSC line (39b) using Q5 Hi-fidelity DNA polymerase and cloned into pBR322. Sanger sequencing of cloned donor showed the BP-elevating rs1173771 haplotype. The BP-lowering rs1173771 haplotype was then constructed by site directed mutagenesis at the 11 SNPs locations in the haplotype using QuikChange II Site-directed mutagenesis kit (Agilent) and confirmed by Sanger sequencing. Isogenic iPSC lines with the precise reconstitution of homozygous BP-elevating rs1173771 and BP-lowering rs1173771 haplotypes were selected, tested, and assessed for quality as described above (**Suppl. Table S6**).

### Differentiation of edited iPSCs to iECs and iVSMCs

The differentiation was performed as we described (**Suppl. Fig. S12**).^31,32^ Briefly, iPSCs were cultured on hESC-qualified Matrigel (Corning) coated 6-cm dishes with mTeSR™ plus medium (STEMCELL Technologies). For the differentiation, iPSCs were dissociated using Accutase (STEMCELL Technologies) and plated at a density of 45,000-50,000 cells/cm^2^ in mTeSR™ plus medium with 10 µM Rock inhibitor Y-27632 (STEMCELL Technologies) on 6-well plates coated with Matrigel. After 24 h, cells were treated with N2B27 medium (a 1:1 mixture of DMEM:F12 with Glutamax and Neurobasal media supplemented with N2 supplement and B27 supplement minus vitamin A; all Life Technologies) plus 8 µM CHIR99021 (Selleck Chemicals) and 25 ng/ml BMP4 (PeproTech) for 3 days to generate mesoderm cells. For EC induction cells were further induced with StemPro-34 SFM medium (STEMCELL Technologies) supplemented with 200 ng/ml VEGF (PeproTech) and 2 µM forskolin (Abcam) for 2 days and purified with CD144 (VE-Cadherin) magnetic beads (Miltenyi Biotec) on day 6. CD144-positive cells were cultured in StemPro-34 SFM medium supplemented with 50 ng/ml VEGF for 6 more days before harvest. For VSMC induction, mesoderm cells were grown with N2B27 medium supplemented with 10 ng/ml PDGF-BB (PeproTech) and 2 ng/ml Activin A (PeproTech) for 2 days. Contractile VSMCs were then induced with N2B27 supplemented with 2 ng/ml Activin A and 2 µg/ml Heparin (STEMCELL Technologies) for 5 days. VSMCs were enriched by removing CD144 + cells using CD144 magnetic beads.

### Immunofluorescence staining for markers of iPSC, iEC, and iVSMC

Cells were grown on matrigel coated coverslips (14mm diameter, Chemglass Life Sciences). Cells were fixed with 4% (vol/vol) paraformaldehyde solution, washed with PBS, permeabilized with 0.25% Triton X-100 (vol/vol) prepared in PBS, for 15 min. For blocking cells were incubated with 10% donkey serum for 45 min at room temperature. This was followed by incubation with relevant antibodies for 1 h at room temperature. For iPSCs characterization, OCT-4 (1:200, AF1759, R&D Systems), NANOG (1:200,ab80892, abcam), TRA-1 (1:100, sc-21706, Santa Cruz) and SSEA4 (1:100, sc-21704, Santa Cruz) primary antibodies were used. For endothelial cell markers, cells were incubated with the anti-human vWF (1:100, A0082, Dako) or VE-cadherin (1:200, AF938, R&D Systems) primary antibodies. For vascular smooth muscle cells, anti-human α-SMA (1:200, sc-130616, Santa Cruz) or Myosin IIb (1:200, 8824S, Cell Signaling Technology) primary antibodies were used. After washing three times with PBS-T (PBS with 0.05% Tween-20) cells were incubated with relevant Alexa-Fluor 488 and Alexa-Fluor 594 secondary antibodies (Invitrogen). Cells were mounted on microscope slides using the ProLong™ Diamond Antifade Mountant with DAPI and images were taken using fluorescence microscopy (Keyence BZ-X800).

### Real-time PCR

RNA extraction and quantitative real-time PCR analysis were performed as described.^33^ Primer sequences are shown in **Suppl. Table S8**. Expression levels of mRNAs were normalized to the endogenous control 18S using ΔΔ^Ct^ method.

### Region capture Micro-C assay and data analysis

Region capture Micro-C was performed largely as described.^34,35^ Dovetail® Micro-C kit was used following manufacturer’s User Guide to create Micro-C libraries. Region capture Micro-C libraries were then generated with reagents from Dovetail® targeted enrichment panels. Indexed Micro-C libraries were pooled together to perform region capture Micro-C. The probe panel targeting human chr5: 32,443,700-33,470,100 (hg38) was designed and synthesized by Twist Bioscience. The panel of 80bp probes covers the rs1173771 LD region and promoters of adjacent protein coding genes. The libraries were sequenced using the Illumina NovaSeq sequencers. Dovetail Micro-C data analysis pipeline was used to generate .hic file for visualization with JuiceBox and .cool file for differential chromatin interaction analysis. Briefly, the sequencing reads were trimmed and cleaned to remove adaptor and low-quality reads with Trim Galore. The cleaned paired sequence reads were aligned to human hg38 genome with bwa. The parse, sort, dedup, and split function of pairtools were sequentially used to record valid ligation event, sort the pairsam file, remove PCR duplicates, and generate .pairs and .bam files, respectively. The .bam file was sorted and indexed with samtools sort and samtools index, respectively. A QC report was generated by a Python file provided by the pipeline. The contact map file (.hic) was generated by juicer too pre with the generated mapped.pairs file. The compressed mapped.pair (by gbgzip) was used to generate .cool file at 1kb or 5kb resolution by cooler cload_pairix. HiCompare^36^ was used to statistically compare chromatin interactions between cells with different genotypes.

## ACKNOWLEDGEMENT

This work was supported by National Institutes of Health grants HL149620 and DK129964.

## AUTHOR CONTRIBUTIONS

HX performed experiments related to rat models. MKM performed experiments related to iPSCs. YL performed region-capture Micro-C analysis. PL analyzed RNA-seq data. MG, KU, MAVA, NB, and AAA contributed to rat studies. YL, MG, and RP contributed to iPSC studies. HX, MKM, YL, PL, AWC, QQ, ASG, SR, and CCO contributed to method development, study design, or data interpretation. ML conceived the study, and AMG and ML designed and led the study. HX, MKM, YL, QQ, AMG, and ML drafted the manuscript.

