## Supplementary tables and figures for "Physiological role and mechanisms of action for a long noncoding haplotype region"

[1]Department of Physiology, Medical College of Wisconsin, Milwaukee, WI, USA; [2]Department of Physiology, University of Arizona College of Medicine – Tucson, Tucson, AZ, USA; [3]Department of Medicine, Division of Nephrology, Hypertension, and Renal Transplantation, University of Florida College of Medicine Gainesville, FL, USA; [4]The Jackson Laboratory, Bar Harbor, ME, USA; [5]Versiti Blood Research Institute, Milwaukee, WI, USA; [6]Department of Pediatrics, Division of Hematology, Oncology, and Transplantation, Medical College of Wisconsin, Milwaukee, WI, USA; [7]Department of Cell Biology, Neurobiology, and Anatomy, Medical College of Wisconsin, Milwaukee, WI, USA.

*co-first authors.

[#]co-corresponding authors:

Aron M. Geurts,

Mingyu Liang,

**Supplementary Table S1: Previously reported associations of rs1173771 with pulse pressure, mean arterial pressure, systolic blood pressure and diastolic blood pressure.** Based on *Nat Genet.* 2011 Sep 11;43(10):1005-11. Similar associations have been identified by other GWAS.

| Region | SNP chr: NCBI 36 posn | Study of origin* (r2#) | Coded allele & freq | Stage 1 | | | | Stage 2 | | | | Combined Stage 1 + Stage 2 | | | |
|---|---|---|---|---|---|---|---|---|---|---|---|---|---|---|---|
| | | | | N effective | Beta | SE | P | N effective | Beta | SE | P | N effective | Beta | SE | P |
| **Pulse pressure association** | | | | | | | | | | | | | | | |
| NPR3-C5orf23 | rs1173771 chr5: 32850785 | Ehret et. al. (0.759) | G 0.525 | 73371 | **0.304** | 0.060 | 3.47E-07 | 43737 | **0.230** | 0.077 | 2.72E-03 | 117108 | **0.276** | 0.047 | 4.56E-09 |
| **Mean arterial pressure association** | | | | | | | | | | | | | | | |
| NPR3-C5orf23 | rs1173771 chr5: 32850785 | Wain et. al. / Ehret et. al. | G 0.525 | 73371 | **0.289** | 0.059 | 1.07E-06 | 43737 | **0.271** | 0.081 | 8.58E-04 | 117109 | **0.283** | 0.048 | 3.51E-09 |
| **Systolic blood pressure association** | | | | | | | | | | | | | | | |
| NPR3-C5orf23 | rs1173771 chr5: 32850785 | | G 0.525 | 73338 | **0.578** | 0.089 | 6.62E-11 | 43736 | **0.416** | 0.114 | 2.53E-04 | 117074 | **0.517** | 0.070 | 1.39E-13 |
| **Diastolic blood pressure association** | | | | | | | | | | | | | | | |
| NPR3-C5orf23 | rs1173771 chr5: 32850785 | | G 0.525 | 73354 | **0.246** | 0.056 | 1.16E-05 | 43740 | **0.198** | 0.075 | 8.25E-03 | 117094 | **0.229** | 0.045 | 3.57E-07 |

**Supplementary Table S2. 11 SNPs in the rs1173771 haplotype region and the genotypes of hiPSC lines Y4 and 39b.**

| pos (hg38) in Chr 5 | variant | BP-lowering allele | BP-elevating allele | allele sequence in iPSC Y4 | allele sequence in iPSC 39b |
|---|---|---|---|---|---|
| 32,814,922 | rs1173771 | A | G | G/G | A/G |
| 32,818,967 | rs9292468 | T | C | C/C | T/C |
| | rs79292845 | T | G | G/G | T/G |
| 32,820,447 | rs2193950 | T | C | C/C | T/C |
| 32,821,105 | rs1173770 | T | C | C/C | T/C |
| 32,828,740 | rs7733331 | T | C | C/C | T/C |
| 32,829,869 | rs1177764 | C | G | G/G | C/G |
| 32,830,415 | rs1173727 | T | C | C/C | T/C |
| 32,831,564 | rs13154066 | T | C | C/C | T/C |
| 32,831,833 | rs12656497 | T | C | C/C | T/C |
| 32,832,368 | rs10059884 | C | A | A/A | C/A |

### Supplementary Table S3: QC reports from the region-capture Micro-C analysis.

| | iEC BP-lowering | | iEC BP-elevating | | iVSMC BP-lowering | | iVSMC BP-elevating | |
|---|---|---|---|---|---|---|---|---|
| Total Read Pairs | 21,325,770 | 100.00% | 27,423,496 | 100.00% | 31,974,380 | 100.00% | 35,721,602 | 100.00% |
| Unmapped Read Pairs | 2,272,572 | 10.66% | 3,186,313 | 11.62% | 4,238,972 | 13.26% | 3,731,879 | 10.45% |
| Mapped Read Pairs | 10,025,033 | 47.01% | 12,560,945 | 45.80% | 14,610,337 | 45.69% | 17,367,970 | 48.62% |
| PCR Dup Read Pairs | 5,425,447 | 25.44% | 5,697,983 | 20.78% | 7,376,485 | 23.07% | 9,237,475 | 25.86% |
| No-Dup Read Pairs | 4,599,586 | 21.57% | 6,862,962 | 25.03% | 7,233,852 | 22.62% | 8,130,495 | 22.76% |
| No-Dup Cis Read Pairs | 3,735,958 | 81.22% | 5,469,880 | 79.70% | 5,842,973 | 80.77% | 6,673,334 | 82.08% |
| No-Dup Trans Read Pairs | 863,628 | 18.78% | 1,393,082 | 20.30% | 1,390,879 | 19.23% | 1,457,161 | 17.92% |
| No-Dup Valid Read Pairs (cis >= 1kb + trans) | 4,175,761 | 90.79% | 6,284,458 | 91.57% | 6,042,987 | 83.54% | 7,475,302 | 91.94% |
| No-Dup Cis Read Pairs < 1kb | 423,825 | 9.21% | 578,504 | 8.43% | 1,190,865 | 16.46% | 655,193 | 8.06% |
| No-Dup Cis Read Pairs >= 1kb | 3,312,133 | 72.01% | 4,891,376 | 71.27% | 4,652,108 | 64.31% | 6,018,141 | 74.02% |
| No-Dup Cis Read Pairs >= 10kb | 2,833,084 | 61.59% | 4,197,159 | 61.16% | 3,958,280 | 54.72% | 5,066,680 | 62.32% |

**Supplementary Table S4: Differential chromatin interactions involving the rs1173771 haplotype region and NPR3 promoter region in iECs or iVSMCs.** IF1 and IF2, chromatin interaction frequencies in cells with BP-lowering and BP-elevating rs1173771 haplotypes, respectively. The rs1173771 haplotype region is at chr5: 32814922-32832368 (hg38). The NPR3 promoter region is at chr5: 32709321-32711521.

| cell type | region1 | start1 | end1 | region2 | start2 | end2 | IF1 | IF2 | D | M | adj.IF1 | adj.IF2 | adj.M | mc | A | Z | p.adj |
|---|---|---|---|---|---|---|---|---|---|---|---|---|---|---|---|---|---|
| iEC | chr5 | 32710000 | 32715000 | chr5 | 32810000 | 32815000 | 2 | 15.40 | 20 | 2.94 | 1.75 | 17.56 | 3.32 | -0.38 | 9.66 | 3.84 | 0.00098 |
| iEC | chr5 | 32705000 | 32710000 | chr5 | 32810000 | 32815000 | 2 | 9.80 | 21 | 2.29 | 1.76 | 11.16 | 2.67 | -0.38 | 6.46 | 3.05 | 0.00922 |
| iVSMC | chr5 | 32709000 | 32710000 | chr5 | 32818000 | 32819000 | 0 | 3.50 | 109 | 2.17 | 1.04 | 4.33 | 2.06 | 0.12 | 2.68 | 2.24 | 0.02487 |

**Supplementary Table S5: Sequences of synthetic sgRNAs for rs1173771 single SNP editing.**

| | **sgRNA sequence** |
|---|---|
| **For SNP rs1173771** | |
| Step 1 sgRNA 1 | 5'-AGUGACUCUCCUCUAAUACA-3' |
| Step 1 sgRNA 2 | 5'-CAUGAGAAUAGCACGGAGAA-3' |
| Step 2 sgRNA 1 | 5'-CAUGAGAAUAGCACGAUUAG-3' |

### Supplementary Table S6: Primer sequences for PCR, cloning, and sequencing.

| Genes | Forward primer sequence | Reverse primer sequence |
|---|---|---|
| ***SNPs region amplification and sequencing*** | | |
| rs1173771 | 5'-CACAGCAAAGTACATAGCAAGGA-3' | 5'-TGATAAACCCATCAGATCTTGTGAG-3' |
| rs9292468 | 5'-GATTGCATTCTTCTTTGGCTGAG-3' | 5'-ACCTATCTGGCTGTCTGTCC-3' |
| rs79292845/ rs2193950 | 5'-GGACCACGGGACATAAACTC-3' | 5'-GGAACATTAGGAGGTACAGATGG-3' |
| rs1173770 | 5'-GTTAGAAGCAAGTTACAAGTAGCAC-3' | 5'-TCTACTGCTATAATTCCAGTTTCCC-3' |
| rs7733331 | 5'-GCCACCAGCTATACTAGAATTACGA-3' | 5'-CTTGCCCATTTATCCCTGAAACC-3' |
| rs1177764 | 5'-AAATGGAGGTTGTTGTGAGGA-3' | 5'-AAAGCATGGAAATGGAAGGGA-3' |
| rs1173727 | 5'-GCAACCTATCCTCTTCCTCAG-3' | 5'-TCACAAATTCTCTAAGTCCTCCC-3' |
| rs13154066/ rs12656497 | 5'-GTTCCCAAAGACACGCTGAG-3' | 5'-CCATAACAACCACCTAAGCCAG-3' |
| rs10059884 | 5'-AGCCACTTCTAAGCCTTATCC-3' | 5'-ACTTCTCTGATCCATAAACCGT-3' |
| ***For LD region deletion confirmation*** | | |
| Across LD region PCR 1 | 5'-AACATTCACCACCAAATTCCAC-3' | 5'-AGAAACTCATGCATTCCGCT-3' |
| Across LD region PCR 2 | 5'-CTCGGTTATTGTGGATAGTGCTG-3' | 5'-GGTCCCTTACAGAAGCAACC-3' |
| ***LD region haplotype donor cloning*** | | |
| LD region haplotype donor amplification | 5'-CACAAGGGTTCCAATTTCTTCAC-3' | 5'-CCACCCTTAGACTCCTTTATCAG-3' |
| LD region haplotype donor cloning in pBR322 | 5'- GAAGAAATTGGAACCCTTGTGCGAGGATGACGATG AGCGC-3' | 5'- TAAAGGAGTCTAAGGGTGGCCTCGCTAACGGATTCACCAC-3' |
| LD region haplotype donor cloning in pBR322 sequencing | 5'-CTCATGTTTGACAGCTTATCATCG-3' | 5'-ACGACAGGAGCACGATCAT-3' |
| ***LD region haplotype reconstruction confirmation*** | | |
| PCR at 5' of LD region | 5'-CTCGGTTATTGTGGATAGTGCTG-3' | 5'-TGATAAACCCATCAGATCTTGTGAG-3' |
| PCR at 3' of LD region | 5'-AGCCACTTCTAAGCCTTATCC-3' | 5'-GGTCCCTTACAGAAGCAACC-3' |
| PCR at 3' of LD region (rs1173770 region) | 5'-GTTAGAAGCAAGTTACAAGTAGCAC-3' | 5'-TCTACTGCTATAATTCCAGTTTCCC-3' |
| ***SNP rs1173771 deletion and re-construction*** | | |
| *For SNP rs1173771 deletion confirmation* | 5'-CAACTTTCAAACGCAGATGGAG-3' | 5'-GTCACCTTTGCTCCAGTTCC-3' |
| *For SNP rs1173771 STEP2 donor PCR* | 5'-AGTCTACTCCCTCCTCATCCT-3' | 5'-CTGTTCCAATCTCTACCTGTTACC-3' |
| *For SNP rs1173771 allele re-construction confirmation* | 5'-AACATTCACCACCAAATTCCAC-3' | 5'-GGAACTGGAGCAAAGGTGAC-3' |

**Supplementary Table S7: Sequences of guide RNAs for rs1173771 haplotype region editing.**

| | Forward gRNA sequence | Reverse gRNA sequence |
|---|---|---|
| **For haplotype rs1173771** | | |
| Step 1 gRNA 1 | 5’-caccgAGTGACTCTCCTCTAATACA-3’ | 5’-aaacTGTATTAGAGGAGAGTCACTc-3’ |
| Step 1 gRNA 2 | 5’-caccgGTCAATGTCGTCAACGCCAA -3’ | 5’-aaacTTGGCGTTGACGACATTGACc -3’ |
| Step 2 gRNA 1 | 5’-caccgAGTGACTCTCCTCTAATGGG-3’ | 5’-aaacCCCATTAGAGGAGAGTCACTc-3’ |

**Supplementary Table S8: Primer sequences for qPCR.**

| Gene | Forward primer sequence | Reverse primer sequence |
|---|---|---|
| Housekeeping and neighboring genes | | |
| 18S | 5'-GATCCATTGGAGGGCAAGTCT-3' | 5'-CCAAGATCCAACTACGAGCTTTT-3' |
| NPR3 | 5'-GGACAGTGAAACCTGAGTTTGAG-3' | 5'-TCATGTAGAGCCAAGACGTAGAG-3' |
| SUB1 | 5'-TCATCTTCTAAACAGAGCAGCAG -3' | 5'-CTGGTTTCATTTCACCTTCAGGA-3' |
| TARS | 5'-CAGTGATTGTTCATCGAGCCA-3' | 5'-CATATTCATCACAGGTTGGTCCC-3' |
| MRPS30 | 5'-GCGAGGTCATATCTTTGCCC-3' | 5'-GAATAATTTCTTCACCACGCACC-3' |

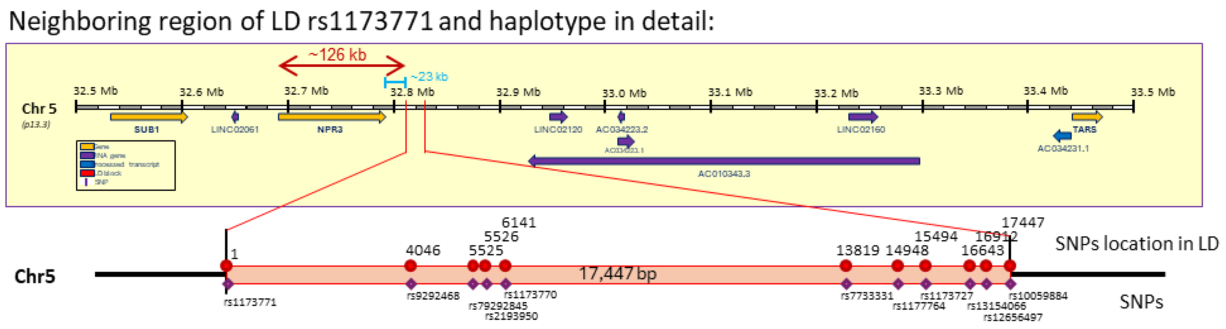

**Supplementary Figure S1: Genomic organization of the rs1173771 haplotype region.** The haplotype contains 11 SNPs in linkage disequilibrium (LD) and spans 17.4 kbp on Chr. 5. The transcription start site of the closest protein-coding gene, *NPR3*, is approximately 126 kbp from the haplotype region.

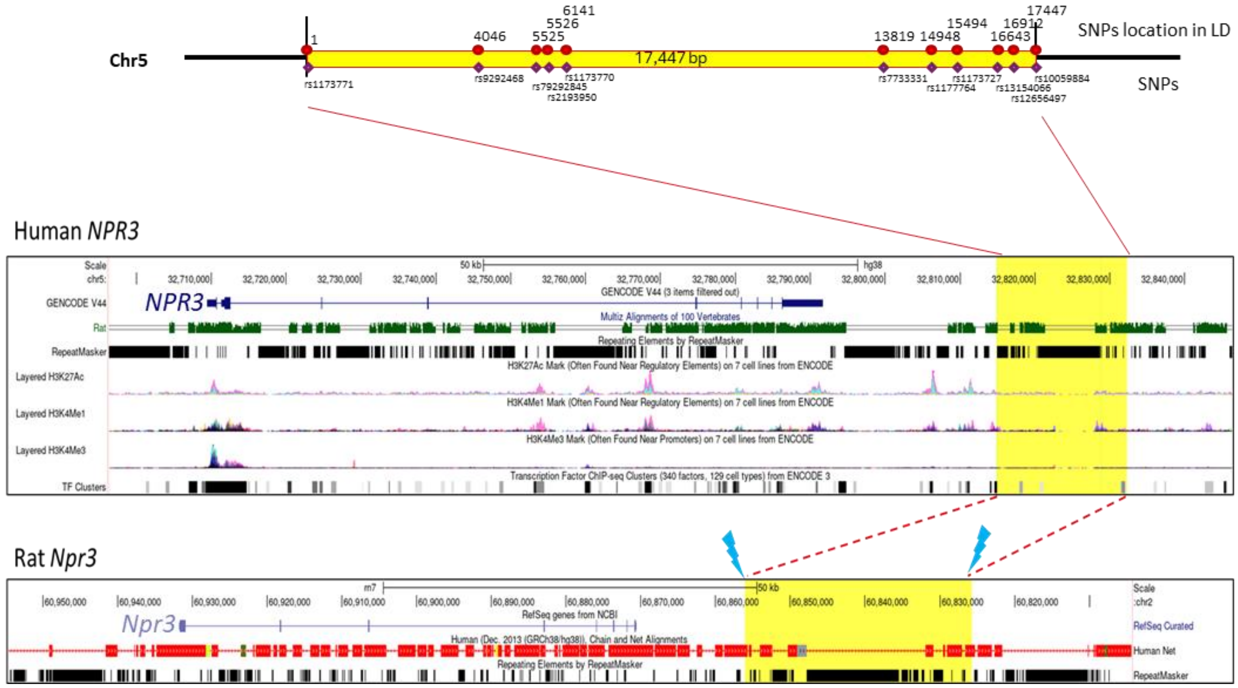

**Supplementary Figure S2. Comparative mapping between the rs1173771 haplotype region in the human genome and the orthologous region in the rat genome.** LiftOver of the ~17.4-kbp region of the human genome identified a ~30.4-kbp region distal to the rat *Npr3 gene*. The region between the blue lightning bolts was deleted using CRISPR-SpCas9. Differences in length appear to be due mainly to repeat expansions within the rat genome compared to human. Chain alignments between the species are shown.

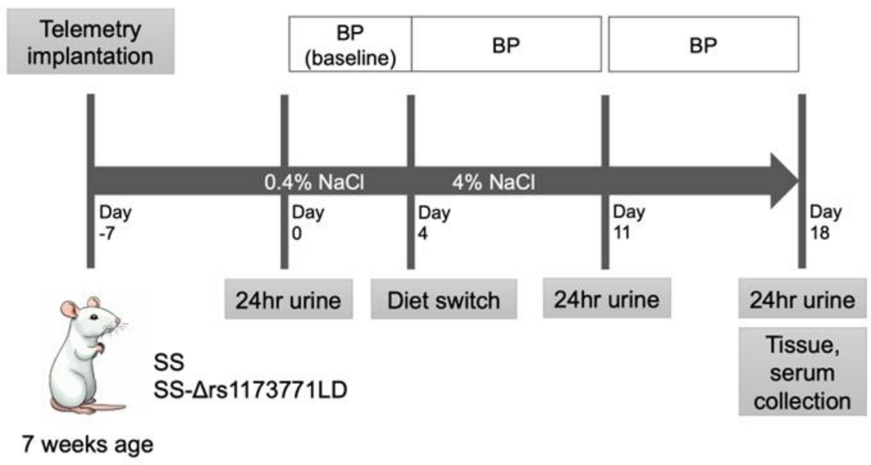

**Supplementary Figure S3. Phenotyping protocol.** BP, blood pressure; SS, Dahl salt-sensitive rat; SS-Δrs1173771LD, SS rats in which a 30.4-kbp noncoding genome segment orthologous to the 17.4 kbp human rs1173771 linkage disequilibrium (LD) region was deleted.

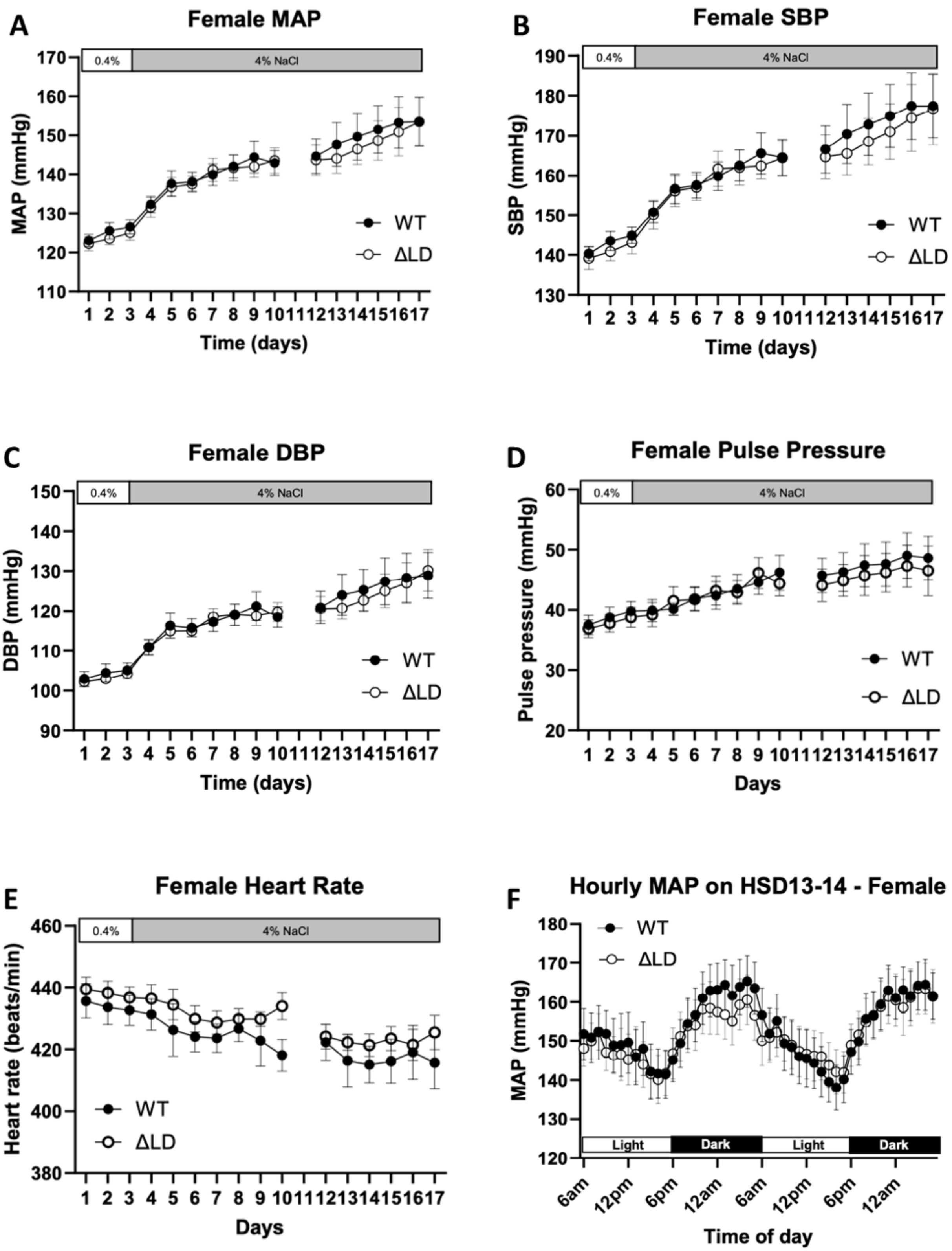

**Supplementary Figure S4. Blood pressure in female rats.** Daily averages of mean arterial pressure (MAP) (A), systolic blood pressure (SBP) (B), diastolic blood pressure (DBP) (C), pulse pressure (D), and heart rate (E) and hourly averages of MAP on day 13-14 of 4% NaCl high-salt diet (HSD) (F) of female SS-Δrs1173771LD$^{-/-}$ rats (ΔLD) and SS littermates (WT) are shown. n=10.

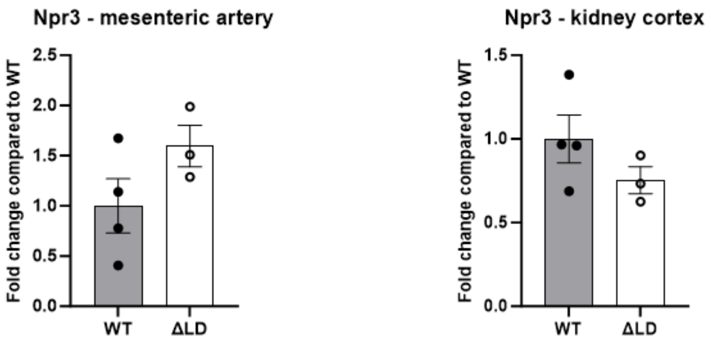

**Supplementary Figure S5. Expression of Npr3 in the mesentery artery and the renal cortex in female SS rats with deletion of the rs1173771 LD orthologous region.** RNA-seq data from female SS-Δrs1173771LD-/- rats (ΔLD) compared with SS littermates (WT) is shown. N=3-4.

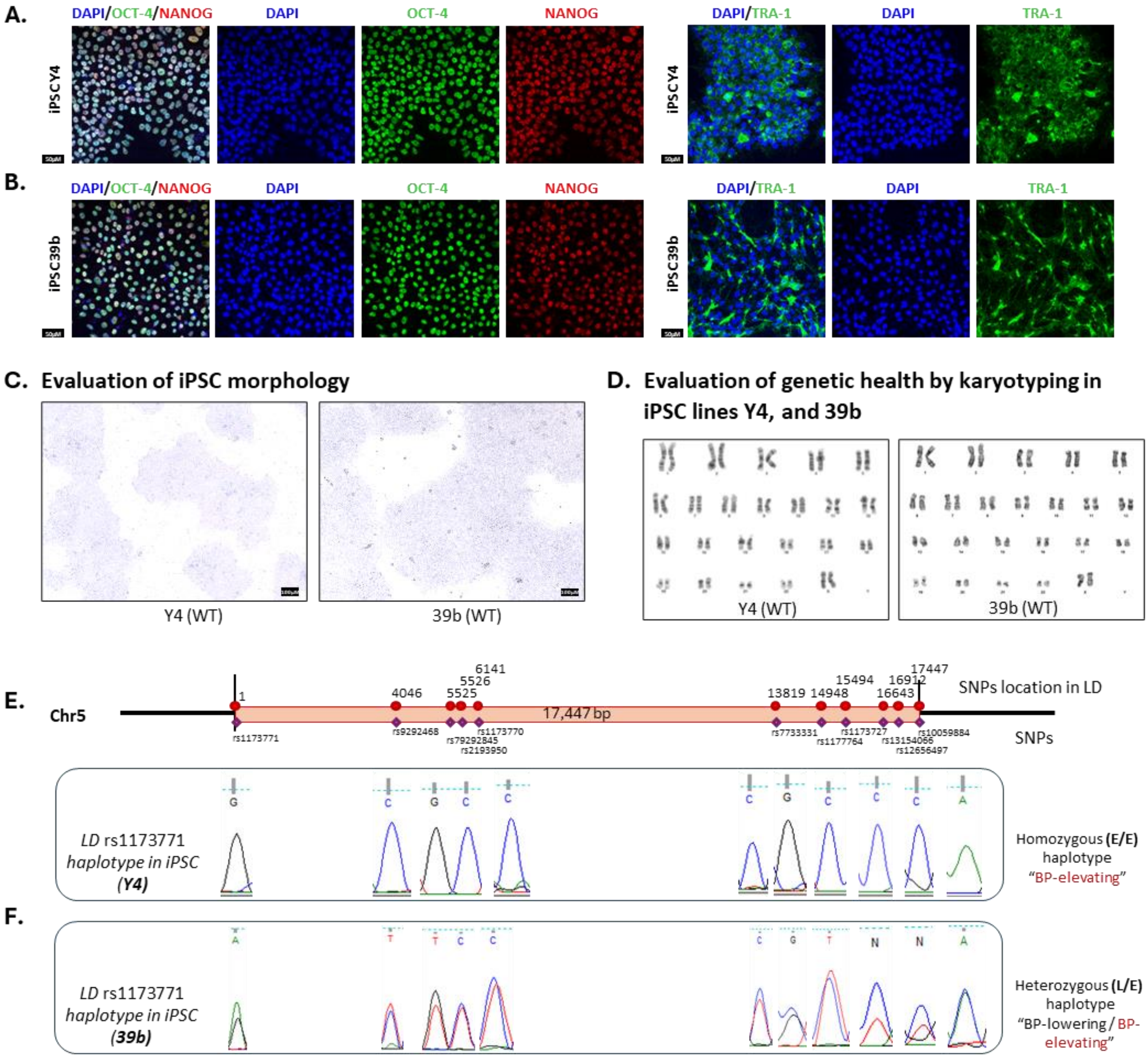

**Supplementary Figure S6: HiPSC lines Y4 and 39b reprogrammed from urine cells.** iPSC lines Y4 **(A)** and 39b **(B)** immunostained for pluripotency markers OCT-4, NANOG and TRA1. iPSC Y4 and 39b were evaluated for morphology **(C)** and karyotype **(D)**. Y4 contains homozygous BP-elevating haplotype at the rs1173771 locus based on Sanger sequencing (**E**). 39b contains heterozygous haplotype at this locus (**F**).

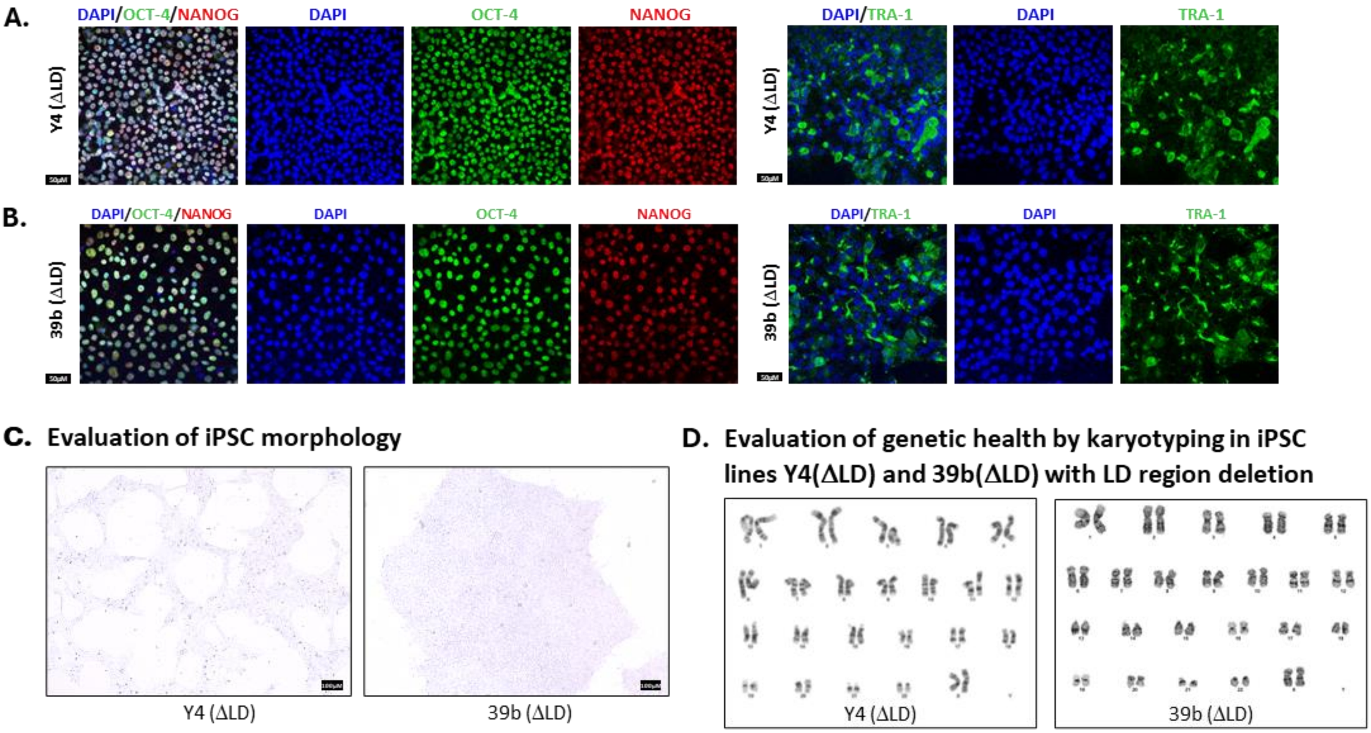

**Supplementary Figure S7: Quality assessment of edited iPSC lines Y4 (ΔLD) and 39b (ΔLD).** Immunostaining for pluripotency markers OCT-4, NANOG and TRA1 (**A, B**), morphology (**C**), and karyotype (**D**) in Y4 (ΔLD) and 39b (ΔLD).

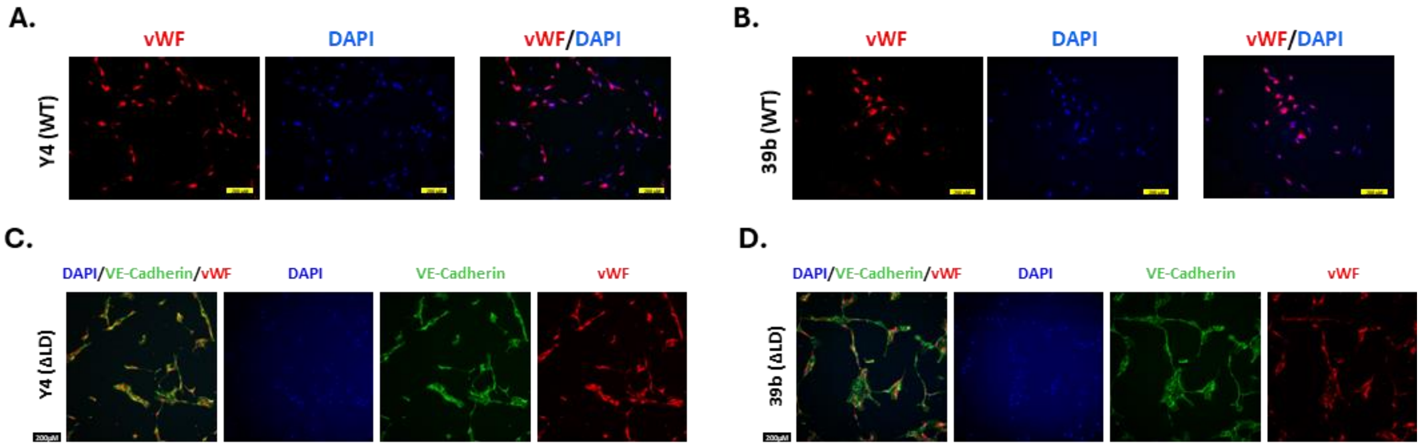

**Supplementary Figure S8**: **Differentiation of Y4, Y4 (ΔLD), 39b, and 39b (ΔLD) to iECs.** Confirmation of iPSC differentiation to iECs by staining of the endothelial cell marker von Willebrand factor (vWF) in iECs derived from wild-type Y4 (**A**) and 39b (**B**) and vWF and VE-Cadherin in iECs derived from Y4 (ΔLD) **(C)** and 39b (ΔLD) **(D).**

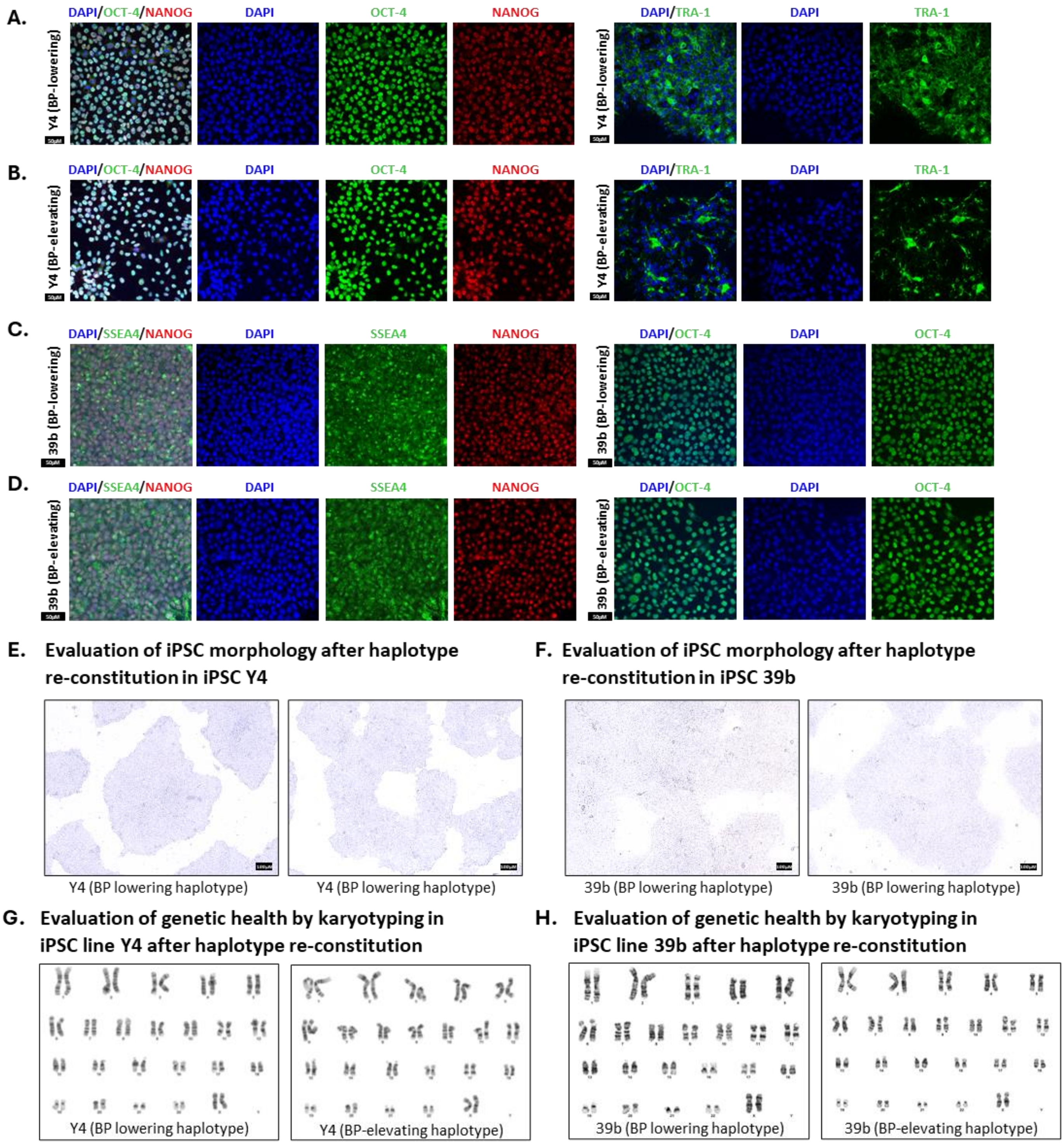

**Supplementary Figure S9: Quality assessment of iPSCs with reconstituted homozygous BP-elevating or –lowering rs1173771 haplotype.** Immunostaining for pluripotency markers OCT-4, NANOG and TRA1 (**A-D**), morphology (**E, F**), and karyotype (**G, H**) in Y4 and 39b cells with reconstituted, homozygous rs1173771 haplotypes.

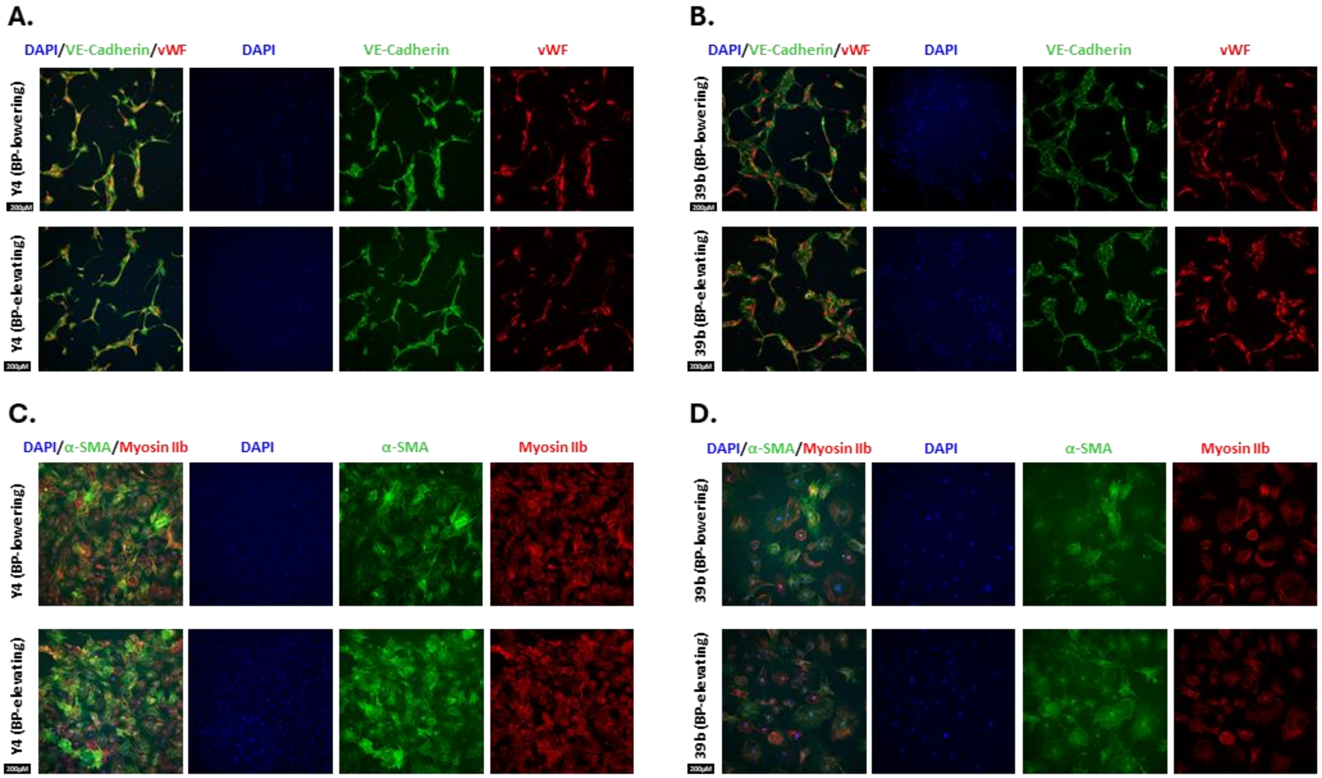

**Supplementary Figure S10: Differentiation of iPSCs with reconstituted homozygous BP-elevating or -lowering rs1173771 haplotype.** Confirmation of iPSC differentiation to iECs shown by immunostaining of an endothelial cell markers VE-Cadherin and von Willebrand factor (vWF) in iECs differentiated from Y4 **(A)** and 39b **(B)** with reconstituted haplotypes. Confirmation of iPSC differentiation to iVSMCs shown by immunostaining of vascular smooth muscle cell markers αSMA and Myosin-IIb in iVSMCs differentiated from Y4 **(C)** and 39b **(D)** with reconstituted haplotypes.

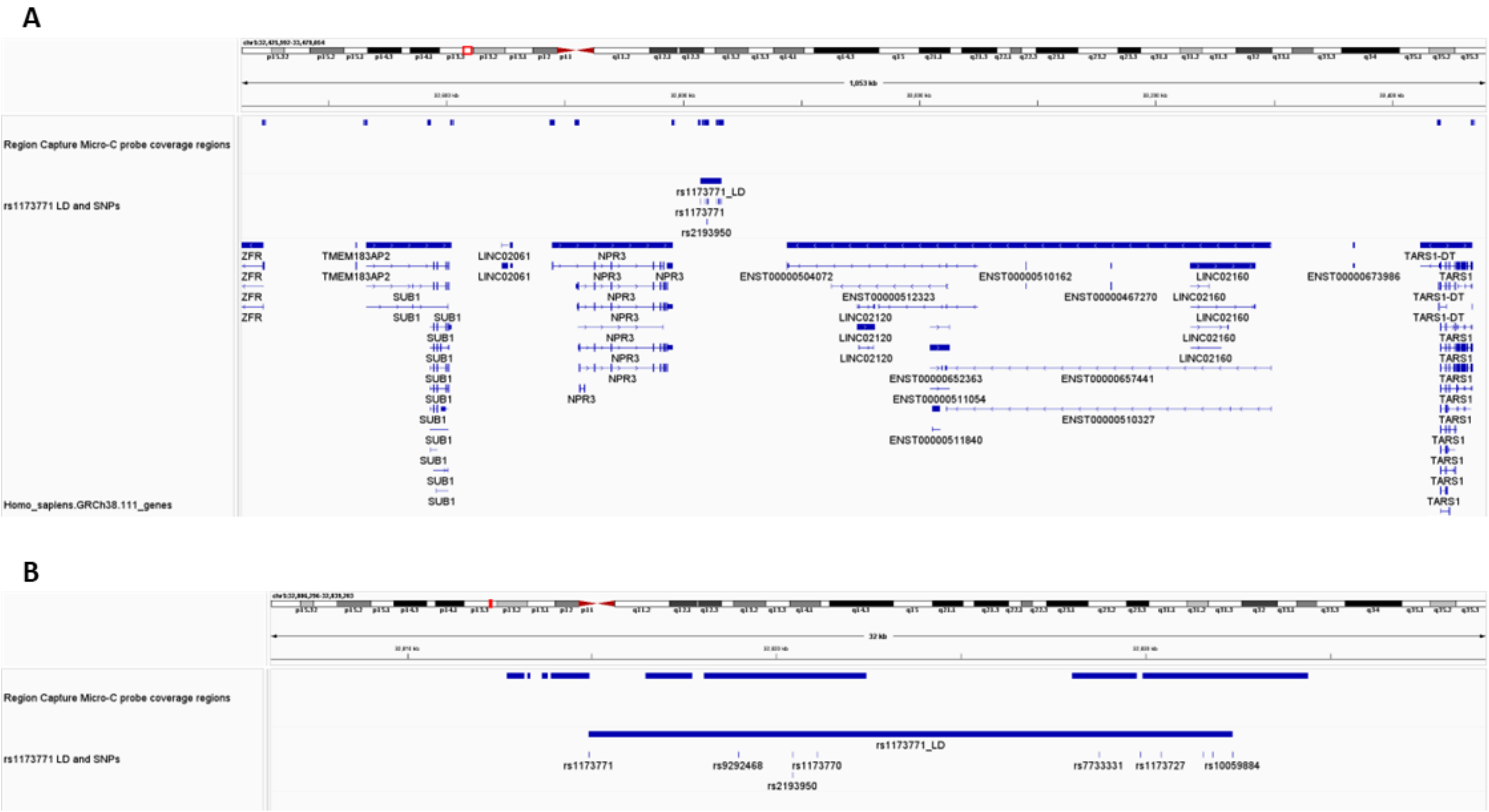

**Supplementary Figure S11: Probe panel targeting human chr5: 32,443,700-33,470,100 (hg38) for region capture Micro-C analysis. A.** The panel of 80bp probes covers the rs1173771 haplotype region and promoters of adjacent protein coding genes. **B.** A zoom-in view of the probe coverage for the rs1173771 haplotype region.

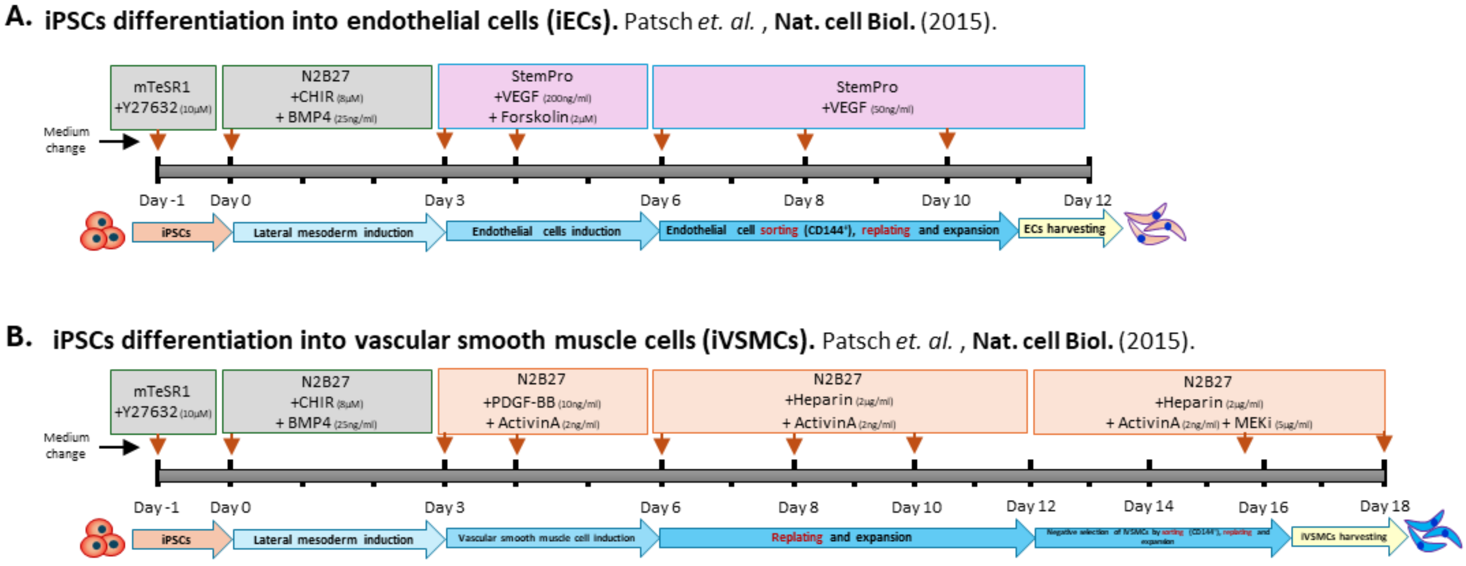

**Supplementary Figure S12: Schematics showing the differentiation protocols for iECs (A) and iVSMCs (B).**
